# Subcellular mRNA localization and local translation of *Arhgap11a* in radial glial cells regulates cortical development

**DOI:** 10.1101/2020.07.30.229724

**Authors:** Louis-Jan Pilaz, Kaumudi Joshi, Jing Liu, Yuji Tsunekawa, Fernando C. Alsina, Sahil Sethi, Ikuo K. Suzuki, Pierre Vanderhaeghen, Franck Polleux, Debra L. Silver

## Abstract

mRNA localization and local translation enable exquisite spatial and temporal control of gene expression, particularly in highly polarized and elongated cells. These features are especially prominent in radial glial cells (RGCs), which serve as neural and glial precursors of the developing cerebral cortex, and scaffolds for migrating neurons. Yet the mechanisms by which distinct sub-cellular compartments of RGCs accomplish their diverse functions are poorly understood. Here, we demonstrate that subcellular RNA localization and translation of the RhoGAP Arhgap11a controls RGC morphology and mediates cortical cytoarchitecture. *Arhgap11a* mRNA and protein exhibit conserved localization to RGC basal structures in mice and humans, conferred by a 5′UTR cis-element. Proper RGC morphology relies upon active *Arhgap11a* mRNA transport and localization to basal structures, where ARHGAP11A is locally synthesized. Thus, RhoA activity is spatially and acutely activated via local translation in RGCs to promote neuron positioning and cortical cytoarchitecture. Altogether, our study demonstrates that mRNA localization and local translation mediate compartmentalization of neural progenitor functions to control brain development.

**Highlights:**

- Arhgap11a in radial glia non-cell autonomously promotes neuronal migration and lamination
- *Arhgap11a* mRNA localizes to radial glial endfeet via a 5’ UTR cis element
- ARHGAP11A expression in basal process and endfeet depends upon its localized mRNA
- Localized mRNA and RhoA-GAP activity in endfeet control radial glial morphology

## Introduction

In eukaryotes, subcellular RNA localization and local translation allow cells to temporally and spatially control functions that rely on dynamic and complex proteomes. In highly polarized cells, such as neurons and migrating fibroblasts, mRNA localization plays a pivotal role in local cytoskeletal regulation and hence local morphology (Buxbaum et al., 2015). Notably, the developing and adult brain contains some of the most highly polarized and elongated cell types found in animals. These features are especially prominent in radial glial cells (RGCs), which control cortical development by acting as neural stem cells to generate neurons and then glia, and by directing neuronal migration (Belvindrah et al., 2007; Malatesta et al., 2000; Mizutani et al., 2007; Noctor et al., 2001; Rakic, 1972; Seuntjens et al., 2009; Siegenthaler et al., 2009; Silva et al., 2019).

RGCs have a unique and highly dynamic bipolar morphology. RGC basal processes emanate from cell bodies in the ventricular zone (VZ) and radially traverse the cortex to form basal endfeet at the pia (Rakic, 2003; Yokota et al., 2010). The basal process can be extremely long, reaching several hundred microns in the mouse and millimeters in humans. The apical and basal endfeet encounter unique niches, with the latter composed of interneurons, Cajal-Retzius neurons and excitatory neurons. Basal endfeet are tightly connected to a basement membrane, forming a barrier between the brain and the overlying meninges, composed of fibroblasts and blood vessels (DeSisto et al., 2020; Haubst et al., 2006; Myshrall et al., 2012). Along the basal process and at endfeet, dynamic filopodia-like extensions extend and retract, which may influence signaling and neuronal migration (Yokota et al., 2010). Further, as development proceeds, basal endfeet become notably more complex in number (Lu et al., 2015) and basal structures are especially complex in ferrets, non-human primates, and humans (Kalebic et al., 2019; Rakic, 1972; Reillo et al., 2017). In line with this, disruptions to RGC morphology can have disastrous consequences on the architecture of the mature cortex by causing cobblestone malformation and lissencephaly (Lambert de Rouvroit and Goffinet, 2001; Myshrall et al., 2012). Altogether this indicates that RGC morphology and subcellular compartmentalization are central to cortical development. Yet, we know surprisingly little about the cellular and molecular mechanisms mediating morphology and function within RGC basal structures.

RGC endfeet are enriched for transcripts encoding cytoskeletal and signaling regulators, including GTPase regulators (Pilaz et al., 2016a; Tsunekawa et al., 2012). Notably, the ubiquitous Rho GTPase is essential for cortical development (Cappello et al., 2012), but whether and how modulation of localized Rho activity controls RGCs is unknown. *Arhgap11a* encodes a RhoA-specific GAP, which promotes GTP hydrolysis and therefore inactivates the small GTPase RhoA (Müller et al., 2018). Arhgap11a has essential roles in modulating the cytoskeleton, including mediating cytokinesis (Zanin et al., 2013), cell invasion (Dai et al., 2018; Kagawa et al., 2013), and neurite outgrowth (Xu et al., 2013). Local synthesis of a Rho regulator, such as Arhgap11a, may thus help dictate subcellular morphology and function of RGCs. However, the requirement of local translation of any transcript in RGCs is unknown.

In this study, we ask whether mRNA localization and local translation of the RhoGAP Arhgap11a mediates RGC morphology and cortical development (**Figure 1A**). We show that both *Arhgap11a* mRNA and protein localize to RGC basal endfeet and processes during cortical development in mice and humans. *Arhgap11a* depletion from RGCs simplifies both basal process and endfeet branching. Acute *Arhgap11a* knockdown in RGCs non-cell autonomously disrupts migration of both excitatory and inhibitory neurons, thus affecting laminar cortical architecture. Using live imaging in mouse tissue we show that the *Arhgap11a* 5’UTR is critical for the active transport of *Arhgap11a* mRNA in the basal process and for its local translation in basal endfeet. Importantly, the endfeet branching phenotype is only rescued when *Arhgap11a* localization is targeted to endfeet via the 5’UTR and when RhoGAP activity is intact. Altogether, our study establishes critical *in vivo* functions of subcellular mRNA localization in neural stem cells of the developing brain.

**Figure 1.**
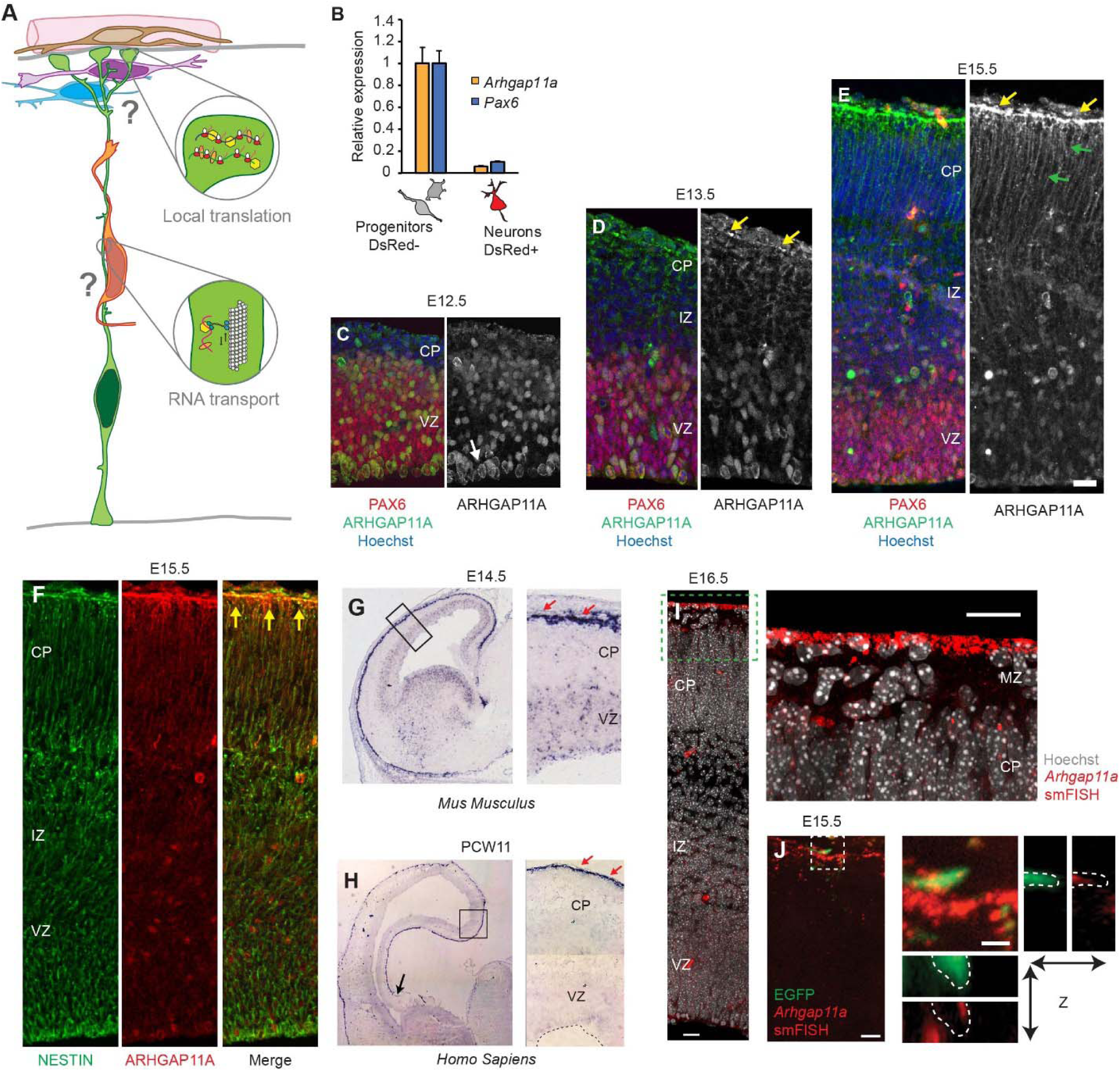
Subcellular localization of Arhgap11a mRNA and protein to RGC basal processes and endfeet during cortical development. (A) Cartoon of a radial glial progenitor with mRNA transport along the basal process and local translation in endfeet. Question marks reflect goal of the present study: what is the role of mRNA subcellular localization and translation in RGCs? (B) qPCR analyses of *Arhgap11a* mRNA levels in E14.5 sorted embryonic cortical cells show *Arhgap11a* expression is specific to neural progenitors. (n=4 embryonic brains, 3 technical replicates). (C-E) Immunofluorescence of ARHGAP11A (green) and RGC cell bodies (PAX6, red). ARHGAP11A localizes to the cell bodies of RGCs at E12.5 (white arrows) (C), additionally begins to localize to RGC basal endfeet (yellow arrows) at E13.5 (D), and at E15.5 is especially enriched in basal endfeet (yellow arrows) and basal processes (green arrows) (E). (F) Immunofluorescence at E15.5, showing ARHGAP11A (red) expression in NESTIN positive RGCs (green). Overlap (yellow signal) is prominent in basal process and endfeet close to the pial surface (yellow arrows). (G,H) *In situ* hybridization of *Arhgap11a* mRNA (purple signal), showing strong enrichment at the pia where RGC basal endfeet reside (red arrows) at E14.5 ((G) and in GW11 human fetal brains (H). (I, J) smFISH *in situ* hybridization depicting *Arhgap11a* mRNA (red) at the pia at E15.5 (I) and in E15.5 brains electroporated with EGFP plasmids confirming the presence of *Arhgap11a* mRNA in RGC basal endfeet (J). In both (I and J), right panels show magnified views of areas highlighted in the left panels. In (J) bottom images show maximum intensity projections of a z-stack. VZ: ventricular zone, CP: cortical plate, IZ: intermediate zone, smFISH : single molecule fluorescent in situ hybridization. Scale bars: C-E, 20μm; I, 20μm; J, left panel: 5μm, right panel: 1μm.

## Results

### *Arhgap11a* mRNA and protein exhibit subcellular localization to RGC endfeet over the course of mouse cortical development

In this study we set out to ask whether the morphology and functions of RGCs are mediated by local translation (**Figure 1A**). From our prior study, we noted that endfeet are enriched for mRNAs encoding cytoskeletal and GTPase signaling regulators (Pilaz et al., 2016a). We therefore sought to investigate functions of transcripts associated with these cellular processes. Arhgap11a stood out as an outstanding candidate, given its established role in cytoskeletal regulation, neurite outgrowth, and GTPase signaling (Kagawa et al., 2013; Müller et al., 2018; Xu et al., 2013; Zanin et al., 2013).

As a first step to examine the role of *Arhgap11a* in RGCs, we assessed its expression during mouse cortical development. Cortical neurogenesis in mice occurs between E11 and E18.5 and in humans, at gestational week (GW) 7-24 (Lodato and Arlotta, 2015; Miller et al., 2019; Silbereis et al., 2016). We used E14.5 *Dcx*-DsRed transgenic mice (Wang et al., 2007) together with FACS and quantitative PCR (qPCR) to quantify *Arhgap11a* mRNA expression in both progenitors (DsRed-negative) and neurons (DsRed-positive) (**Figures 1B,S1A**). *Arhgap11a* was preferentially expressed in dividing progenitors and notably absent from newborn excitatory neurons. Likewise, in single-cell RNA-seq datasets of the developing mouse and human cortex, *Arhgap11a* was highly enriched in RGCs but absent from post-mitotic excitatory and inhibitory neurons (**Figure S1B-D**) (Loo et al., 2019; Nowakowski et al., 2017; Telley et al., 2019). These data demonstrate that *Arhgap11a* expression in the developing cerebral cortex is largely specific to progenitors, including RGCs.

Next, we used immunohistochemistry to evaluate the expression pattern of ARHGAP11A protein in the developing mouse cortex. At E12.5 in the mouse cortex, ARHGAP11A localized within the germinal zones (**Figure 1C**). However, strikingly, beginning at E13.5, ARHGAP11A protein became enriched near the pial surface, at presumptive RGC endfeet and along basal processes (**Figure 1D**). This pial localization of ARHGAP11A was especially enriched at E15.5. when it showed a gradient of expression in the basal process with higher expression nearest to the pia and within the CP, and lower levels in the IZ (**Figure 1E**). We verified ARHGAP11A expression and localization in RGCs by co-staining with the RGC intermediate filament marker NESTIN at E15.5 (**Figure 1F**). This timing of ARHGAP11A localization to basal process and endfeet of RGCs coincides with the onset of increased branching of endfeet (Lu et al., 2015), suggesting that Arhgap11a could function in RGC basal morphology.

We next examined *Arhgap11a* mRNA localization over the course of corticogenesis, to determine if its spatial and temporal localization pattern matched that of the protein. Towards this end, we used traditional as well as single-molecule fluorescent *in situ* hybridization (smFISH). At E14.5, *Arhgap11a* mRNA was detected at the pial surface, at a significantly higher level than in the VZ (**Figure 1G**). Reinforcing the specific expression in RGCs, *Arhgap11a* co-localized with EGFP labeled RGCs, but was notably absent from Tuj1 positive neurons (**Figure S1E**). At E15.5 and E16.5, *Arhgap11a* localization at the pia was especially prominent. smFISH of sections containing EGFP-labeled RGCs confirmed that *Arhgap11a* mRNA is present in RGC basal processes and endfeet (**Figure 1I, J**).

Given this striking localization pattern of *Arhgap11a* to RGC endfeet, and the conserved expression within RGCs of mice and humans, we next tested whether *Arhgap11a* mRNA is also present in the basal endfeet region of the human developing cortex. Thus, we employed traditional *in situ* hybridization performed on brain sections from post-conceptional week 11. This demonstrated that *ARHGAP11A* mRNA exhibits conserved enrichment to the pia in humans (**Figure 1H**), which are in line with a previous report (Florio et al., 2018). Altogether, these data demonstrate that *Arhgap11a* RNA and protein exhibit concordant localization in a developmentally controlled fashion to RGC basal processes and endfeet.

### Arhgap11a controls RGC basal process morphology and non-cell autonomously controls radial migration of excitatory neurons

ARHGAP11A localization to RGC basal processes and endfeet coincides developmentally with increasing morphological complexity of these structures (Lu et al., 2015). Further, excitatory neurons rely on the integrity of RGC basal processes to migrate from the germinal zones to the cortical plate (Belvindrah et al., 2007; Elias et al., 2007; Nakagawa et al., 2019; Noctor et al., 2004). Given that ARHGAP11A has established functions in modulating RhoA signaling and cytoskeletal morphology (Müller et al., 2018; Xu et al., 2013; Zanin et al., 2013), we hypothesized that *Arhgap11a* may regulate basal process morphology and thus influence excitatory neuron migration. To evaluate these possibilities, siRNAs targeting the 3’UTR of *Arhgap11a* mRNA were introduced by *in utero* electroporation together with a membrane-localized EGFP reporter driven by an RGC-specific promoter (pGLAST-EGFP-CAAX, **Figure 2A**). We sought to manipulate *Arhgap11a* expression at E15.5, reasoning that alterations to basal process complexity should be evident at this critical timepoint (Lu et al., 2015). Using siRNAs, we effectively depleted *Arhgap11a* in RGCs both at the mRNA and protein level, in RGC cell bodies and basal endfeet (**Figure 2B-D**, bins 1 and 10, respectively). We observed very little FISH signal in the IZ or CP within the control, thus reinforcing the specific expression of *Arhgap11a* in RGCs but not neurons.

**Figure 2.**
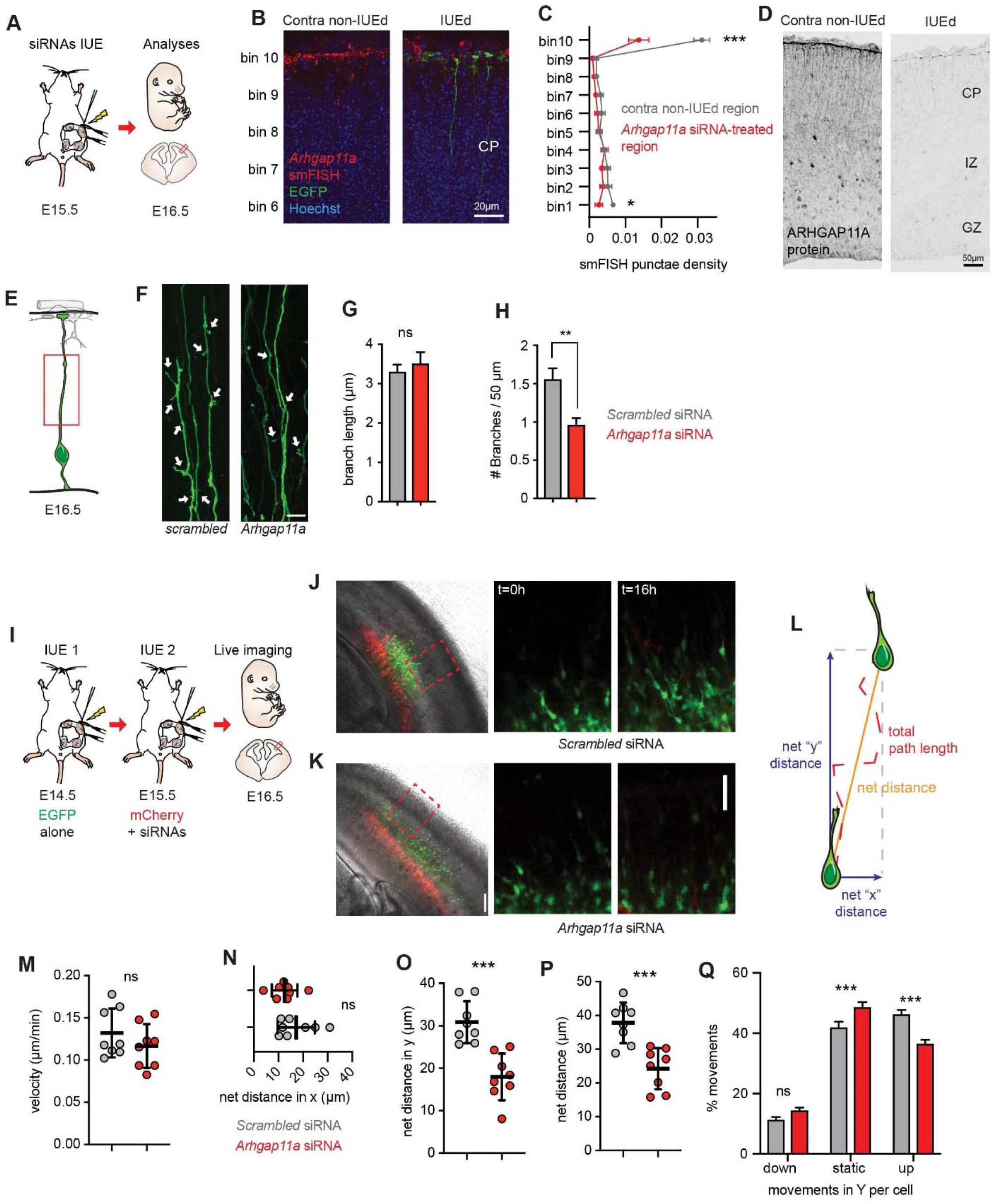
*Arhgap11a* controls RGC basal process morphology and non-cell autonomously controls radial migration of excitatory neurons. (A) Schematic overview of the experiments in (B-H). (B-D) Electroporation of siRNAs targeting *Arhgap11a* efficiently deplete *Arhgap11a* mRNA and protein from the developing cortex. Depletion of *Arhgap11a* mRNA in endfeet in the electroporated region (IUE, green) after siRNA treatment, as evidenced by *Arhgap11a* smFISH (red) and immunofluorescence (D). (C) Binned quantification of *Arhgap11a* smFISH punctae in electroporated and contralateral non-electroporated regions. Bin 1 corresponds to the region lining the ventricle, and Bin 10 to the region adjacent to the meninges. (E) Cartoon of regions analyzed in RGC basal processes. (F) EGFP electroporated RGCs depicting reduced branches (arrows) along the basal process following *Arhgap11a* knockdown. (G) *Arhap11a* depletion in RGCs does not significantly impact the length of branches along the RGC basal process. (scrambled: n=101 branches from 3 brains, 3 independent experiments, *Arhgap11a*: n=72 branches from 3 brains, 3 independent experiments, unpaired t-test with Welch’s correction). (H) *Arhap11a* depletion in RGCs significantly decreases the density of branches along the RGC basal process. (Scrambled: n=112 cells from 6 brains, 5 independent experiments, *Arhgap11a*: n=99 cells from 5 brains, 4 independent experiments, unpaired t-test with Welch’s correction). (I) Schematic overview of the experiments in (J-Q) aimed at testing the impact of *Arhgap11a* depletion in RGCs on neuronal migration. Sequential IUEs were performed to label neurons (EGFP, green) and RGCs (red) at E16.5 when analysis is performed. (J,K) Representative images showing electroporated regions (left panels) and position of migrating of neurons (green) at the beginning (t=0 hrs, middle panels) and end of the live-imaging experiment (t=16 hrs, right panels). (L) Neuronal migration parameters analyzed. (M-P) *Arhgap11a* depletion in RGCs does not significantly affect either velocity of migration (M), nor net-distance in X trajectory (N), but significantly reduces net distance travelled in the Y trajectory (O) and compiled direction (P). (Q) *Arhgap11a* knockdown in RGCs non-cell autonomously causes neurons to undergo more static movements and fewer movements toward the cortical plate (up). (Scrambled and *Arhgap11a*: n=8 brains from 2 independent experiments, unpaired t-tests). siRNAs: small interfering RNAs, IUE: *in utero* electroporation, CP: cortical plate, IZ: intermediate zone. *: p-value<0.05. **: p-value<0.01. ***: p-value<0.001. Scale bars: B: 20μm, D,J-K right panels 50μm, F: 10μm, J-K left panels: 100μm. Bar plots indicate means +/-SEM.

We next examined morphology along the RGC basal process. For this we used 3D reconstructions of either control or siRNA-treated EGFP+ basal processes to quantify the density and length of extensions in the basal process (**Figure 2E-H)**. There was no impact of *Arhgap11a* depletion upon the length of small extensions (average branches were 3.3 μm and 3.5 μm for *scrambled* and *Arhgap11a* conditions, respectively) (**Figure 2G**). However, we noted a significant decrease in the density of these cellular extensions in brains transfected with *Arhgap11a* siRNAs compared to *scrambled* siRNAs (**Figure 2H**). This data demonstrate that *Arhgap11a* regulates branching complexity along the RGC basal process.

The integrity of the RGC basal process is paramount for proper radial migration of excitatory neurons from the ventricular zone to the cortical plate (Belvindrah et al., 2007; Elias et al., 2007). Indeed, ectopic basal process branching can alter patterns of migrating neurons, including speed, directionality, and pausing (Nakagawa et al., 2019). Therefore, we assessed whether aberrant RGC morphology induced by *Arhgap11a* loss could also impact neuron migration in a non-cell-autonomous manner. To do this we used a paradigm relying on sequential *in utero* electroporation on consecutive days (**Figure 2I**). At E14.5 brains were transfected with an EGFP plasmid to label newborn migratory neurons, one day later (E15.5). At E15.5 we then performed an IUE in the identical location using scrambled or *Arhgap11a* siRNAs and membrane-bound mCherry to label the RGC scaffold which these neurons would migrate upon. Importantly, since *Arhgap11a* is not expressed in migrating excitatory neurons (**Figures 1, S1**), this allowed us to quantify migration of WT neurons along an *Arhgap11a* deficient RGC scaffold. At E16.5 we performed overnight (16h) live imaging of EGFP+ migrating neurons along mCherry+RGCs in organotypic brain slices (**Figure 2I-K**). In particular, we focused on migration within the SVZ and IZ where both mCherry+ RGCs and EGFP+ neurons were labeled.

From live imaging, we quantified several parameters related to neuronal migration: average speed, maximum speed, total distance, net distance, net distance in the Y and X axes (in the radial or tangential dimensions, respectively), and fraction of time spent mobile including towards the CP or VZ (**Figure 2L-Q, S2A-D**). Loss of *Arhgap11a* in RGCs had no impact upon average velocity or maximum velocity, total distance, and net distance of neurons traveled in the X axis (**Figure 2M,N, S2C,D**). However, there was a striking and significant decrease in the net migrated distance and a particularly strong decrease in the net distance in the Y axis (**Figure 2O,P**). Additionally, in brains where *Arhgap11a* was depleted in RGCs, radially migrating neurons spent more time immobile (static) and showed decreased propensity to move towards their final destination in the CP (**Figure 2Q**). These findings indicate that *Arhgap11a* loss does not impair the speed or ability of neurons to migrate per se, but clearly reduces the efficiency and trajectory of radial migration. Altogether, these results show that expression of *Arhgap11a* in RGCs non-cell autonomously controls radial migration of neurons along basal processes, linking altered RGC morphology to aberrant neuronal migration.

### Arhgap11a functions non-cell autonomously in RGCs to control laminar organization

Our live imaging experiments demonstrate that acute depletion of *Arhgap11a* from RGCs has a non-cell autonomous impact upon radial migration, begging the question of whether *Arhgap11a* expression in RGCs influences cortical lamination. The six layered laminar structure of the cortex is critical for establishing proper circuitry in the brain. To investigate the impact of acute *Arhgap11a* depletion upon cortical lamination, we used *in utero* electroporation to introduce *Arhgap11a* or scrambled siRNAs in the embryonic brain at E15.5 (**Figure 3A**). Scrambled siRNAs were co-electroporated with CFP while *Arhgap11a* siRNAs were co-electroporated with GFP, enabling us to discriminate experimental conditions in post-natal mice.

**Figure 3.**
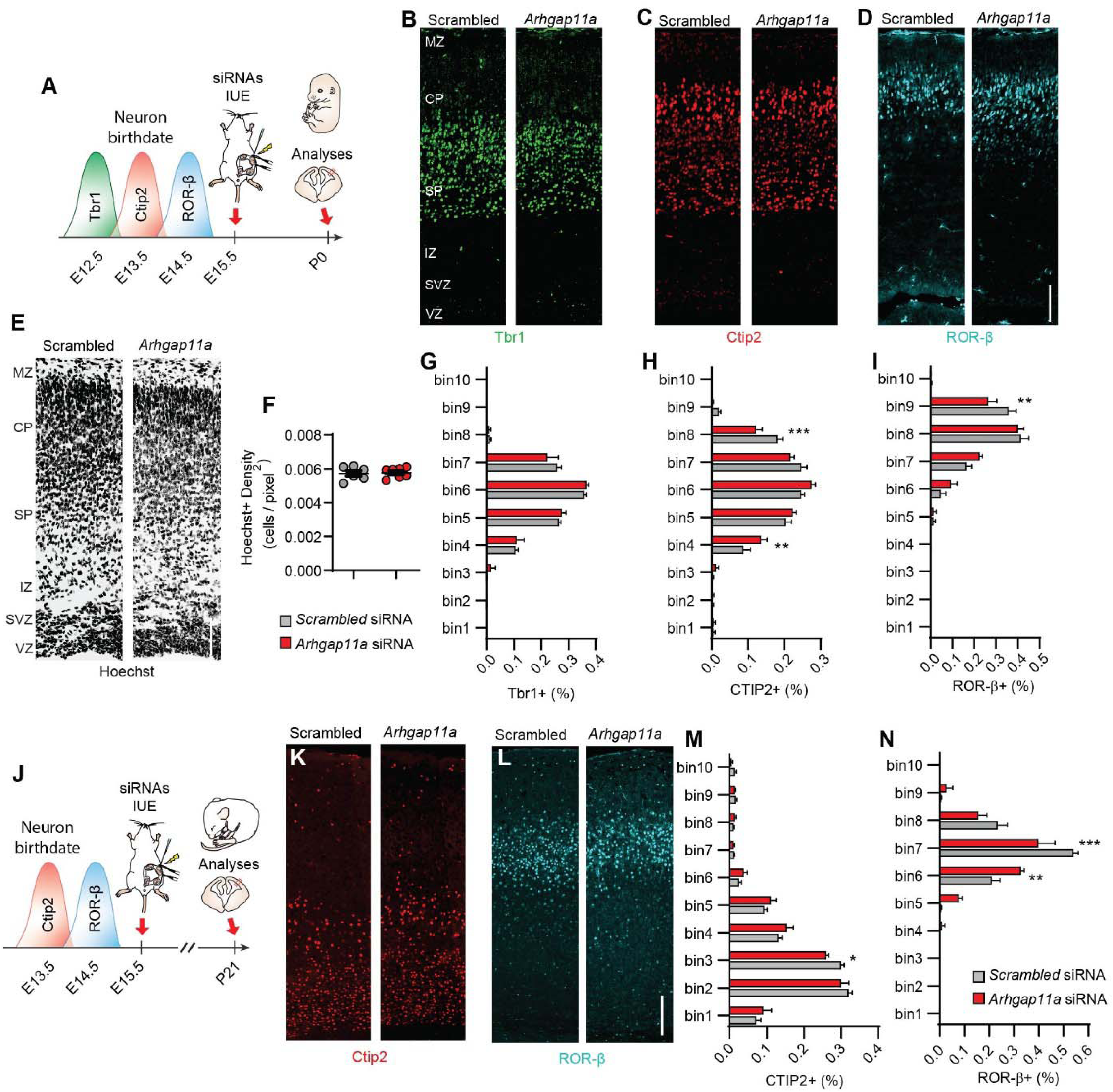
Arhgap11a non-cell autonomously controls laminar architecture. (A) Schematic overview of the experiments in (B-I) which test the impact of acute *Arhgap11a* depletion at E15.5 on neuron positioning at P0. Waves of peak generation of excitatory neuronal subtypes generated prior to electroporation are represented. (B-E) Representative images showing immunofluorescence of Tbr1 (green, B) Ctip2 (red, C), RorB (cyan, D) or Hoechst (E) in P0 brains IUE’d with either scrambled or *Arhgap11a* siRNAs at E15.5. (F) Acute depletion of *Arhgap11a* at E15.5 in RGCs does not impact nuclei density in the cerebral cortex at P0. (Scrambled: n=6 brains from 3 IUE’d brains; Arhgap11a: n=7 brains from 3 IUE’d brains).(G-I) Acute depletion of *Arhgap11a* at E15.5 in RGCs has no impact on the distribution of early-born Tbr1+ neurons (G), but causes fewer later-born Ctip2+ (H) and RorB+ (I) neurons in the upper layers (bins 8, 9) in the cerebral cortex at P0. (Scrambled: n=5-6 brains from 3 independent experiments, *Arhgap11a*: n=6-7 brains from 3 independent experiments. 2-way ANOVA with Bonferroni post-tests).(J) Schematic overview of the experiments in (J-N) which test the impact of acute *Arhgap11a* depletion at E15.5 on neuron positioning at P21. (K-N) Acute *Arhgap11a* depletion in RGCs at E15.5 causes fewer Ctip2+ (M) and Ror-B+ (N) neurons in bin3 and bins 7, respectively, at P21. (Scrambled: n=7 brains from 3 independent experiments, *Arhgap11a*: n=4 brains from 3 independent experiments. 2-way ANOVA with Bonferroni post-tests). siRNAs: small interfering RNAs, IUE: *in utero* electroporation. VZ: ventricular zone, SVZ : subventricular zone, CP: cortical plate, IZ: intermediate zone. *: p-value<0.05, **: p-value<0.01. ***: p-value<0.001. Scale bars: B-E: 200μm, K,L: 100μm.

The fluorescent markers were used to label the IUE’ed region in the mouse cortex. We noted no significant difference in the distribution of labeled cells at P0 in either condition (**Figure S3A-C**). Further, the overall density of cells in the cortex was not affected (**Figure 3E-F**), indicating the generation and survival of neurons were not altered after *Arhgap11a* knockdown.

We then examined laminar organization in the cortex at P0 (**Figure 3A-I**). At this stage, most projection neurons have completed their migration to the CP. We immunostained P0 cortical sections for markers of deep and superficial layer neurons. Using this strategy, we could simultaneously examine the impact of acute *Arhgap11a* reduction upon neurons born several days prior to the siRNA treatment (layers 5/6) as well as with peak genesis one day prior to electroporation (layer 4) (**Figure 3A**). Further, since it takes 24 hours (Greig et al., 2013; Silva et al., 2019) for neurons to complete their migration along RGC basal processes, we predicted that layer IV neurons and possibly layer V neurons may show impaired lamination, whereas layer VI neurons would be unaffected. This strategy is further appropriate given that *Arhgap11a* expression is not detected in neurons.

Tbr1 is highly expressed in early-born neurons of the preplate and layer VI neurons, which arise at E12.5 and migrate into the CP within 24 hours (**Figure 3A, B**). Tbr1+ neuron distribution was similar between knockdown and control mice (**Figure 3B,G**). This is consistent with our prediction that migration of layer VI neurons along the basal process is complete prior to E15.5 siRNA treatment. CTIP2 primarily labels neurons in layers V and VI, whose peak generation occurs around E13.5, two days prior to electroporation (**Figure 3A,C**). This population showed a slight but significant altered distribution in *Arhgap11a*-depleted brains (**Figure 3C,H**). This is consistent with the fact that some layer V, VI neurons are still completing radial migration at the time of acute RGC insult. ROR-ß is expressed primarily in layer IV neurons whose peak birth is evident at E14.5 (**Figure 3A, D**). Compared with scrambled control, the distribution of ROR-ß+ neurons was skewed in the cortex after *Arhgap11a* knockdown in RGCs (**Figures 3D,I**). Consistent with a migration defect, fewer ROR-ß+ neurons localized in superficial layers. These data show that an *Arhgap11a* deficient RGC scaffold is associated with impaired migration and organization of layer IV and V neurons born 1-2 days earlier. This lamination defect is a clear outcome linked to impaired radial migration defects observed with live imaging. Further these results suggest a short-term impact of altered RGC morphology upon cortical lamination.

We next asked whether acute *Arhgap11a* depletion could influence longer-term lamination patterns, by repeating similar experiments and instead collecting brains at P21 (**Figure 3J**). Notably, the placement of Ctip2+ and ROR-ß+ neurons in the *Arhgap11a*-deficient cortex was significantly altered, particularly for layer IV neurons (**Figure 3K-N**). This mild phenotype, nevertheless, is striking in light of the fact that it resulted from an acute embryonic knockdown about 25 days earlier. Altogether, these results indicate that acute *Arhgap11a* knockdown in RGCs dysregulates the laminar organization of cortical neurons at both P0 and P21.

The laminar defects we observed could be caused by cell-autonomous requirements of *Arhgap11a* in neurogenesis of RGCs, which could impact production of Layer 4 neurons. To test this possibility, we thus performed IUE with siRNAs against *Arhgap11a* at E15.5 and quantified proliferation of progenitors in E17.5 brains (**Figure S3D-I**). RGCs and IPs were assayed using Sox2 and Tbr2 staining, respectively. Global proliferation was measured by Ki67 staining. Importantly, there was no difference in the total number of dividing cells or the number of neural progenitors in the germinal zones (i.e., VZ plus SVZ) after depleting *Arhgap11a* for 48hrs (**Figure S3I**). This finding indicates that short-term alterations in progenitor proliferation do not explain the laminar defects.

### *Arhgap11a* controls distal RGC basal process morphology and interneuron numbers in the marginal zone

RGC basal process morphology is dynamic and complex not only in the IZ but also near the pial surface in the marginal zone (MZ) (Lu et al., 2015; Yokota et al., 2010). Indeed, morphological analyses of RGC basal process morphology in MZ across mouse cortical development shows that endfeet number is constant through E14.5, and increases by E16.5 (Lu et al., 2015). Moreover, consistent with observations by others (Yokota et al., 2010) we observed by live imaging in tissue that basal processes can be highly dynamic in the MZ of E16.5 embryonic brain slices (VIDEO S1). Given that ARHGAP11A localization to RGC basal processes and endfeet coincides developmentally with RGC dynamic nature and complexity in the MZ (Lu et al., 2015) (**Figure 1**), we tested whether Arhgap11a also regulates basal process complexity in this basal region.

Towards this, we employed 3D reconstructions of either control or siRNA-treated EGFP+ basal processes, introduced by IUE at E15.5. To assay RGC morphology, we reconstructed the RGC arbor in the MZ and quantified branching points of basal processes within the marginal zone (MZ)(**Figure 4A, B**). A 5μm cut-off was used to designate branches (Wong et al., 2017). In RGCs transfected with scrambled siRNAs (control), we observed multiple types of branches emanating from the basal process in the MZ, as described previously at this stage (**Figure 4C,D**) (Lu et al., 2015; Yokota et al., 2010). In contrast, *Arhgap11a* siRNA knockdown induced a significant decrease in branching complexity of the basal process in the MZ (**Figure 4C,D**). Relative to control, there was also a significant reduction in the total number of branches in *Arhgap11a* depleted RGCs (**Figure 4E**). This effect persisted after 48h, albeit to a lesser extent (**Figure S4A-C**).

**Figure 4.**
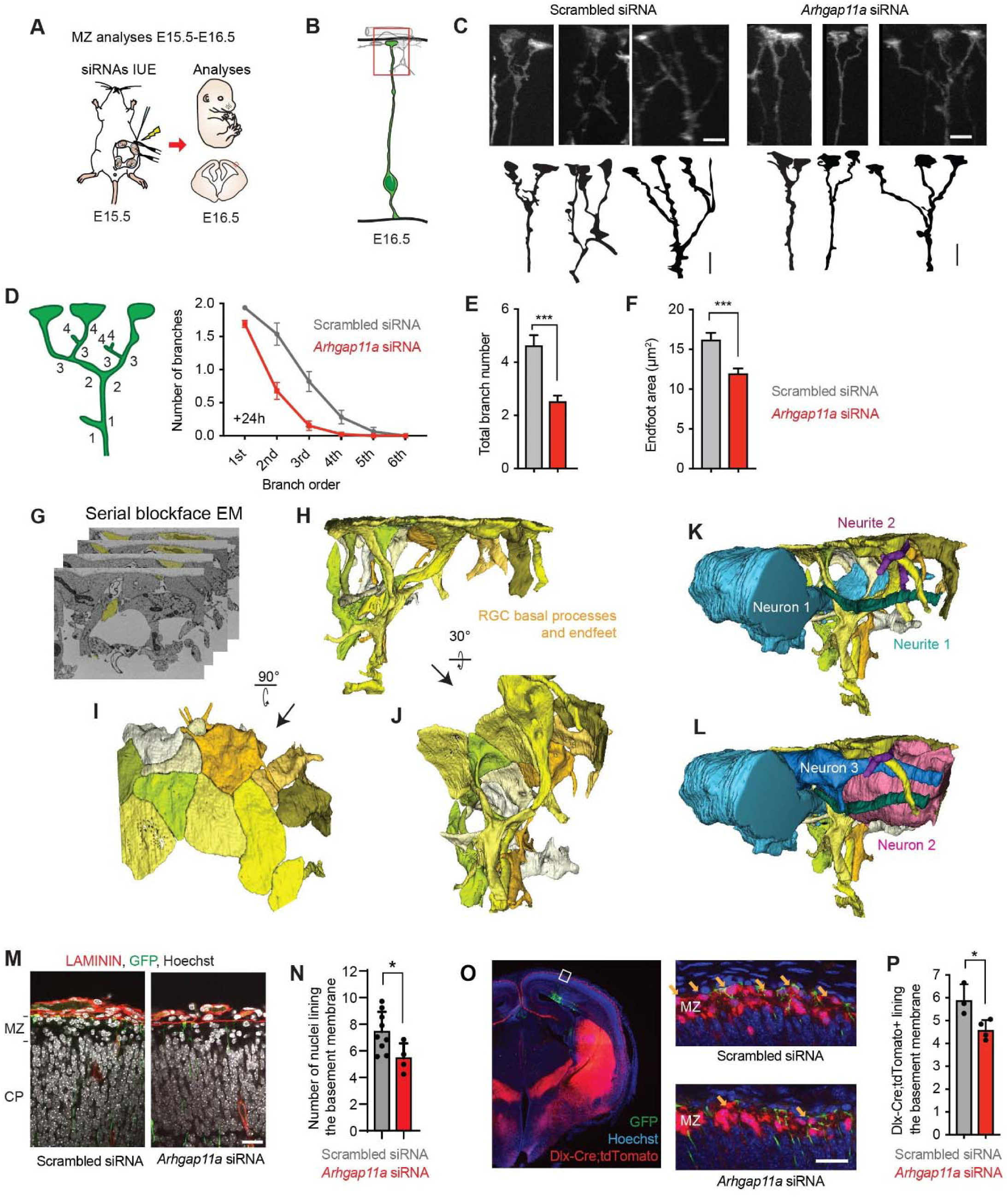
*Arhgap11a* promotes RGC basal process and endfeet complexity and interneuron numbers in the marginal zone. (A) Schematic overview of the experiments in (B-F) which examine acute impact of RGC knockdown upon RGC basal process and endfeet in the marginal zone). (B) Region analyzed in the experiments. (C) Representative images showing basal process and endfoot complexity in the MZ in IUE’d RGCs. A tracing of images is included below. (D-F) Method used to define branch orders in RGC basal processes in MZ (D, left) and quantification of significant reduction in branch complexity (D, right), average total branch number per RGC (E) and area of endfeet at the basement membrane (F). (Scrambled: n=78 cells from 6 brains, 4 independent experiments, *Arhgap11a:* n=78 cells from 5 brains, 4 independent experiments, unpaired t-test). (G-L) 3D reconstructions of the MZ niche using serial blockface electronic microscopy shows tight interactions between presumed (blue, pink) interneurons, cajal retzius neurons, and RGC basal processes and endfeet (yellow). (M) Immunofluorescence depicting nuclei (white, Hoechst), laminin (red) in GFP (green) electroporated regions. (N) Quantification of significant reduction in the number of nuclei lining the basement membrane in *Arhgap11a* depleted brains. (Scrambled: n=9 brains from 5 independent experiments, *Arhgap11a:* n=4 brains from 4 independent experiments, unpaired t-test). (O) Immunofluorescence depicting Hoechst+nuclei (blue), tdTomato+ interneurons (red) in GFP (green) electroporated region, with higher magnification images shown on right. (P) Quantification of significant reduction in Tomato+ interneurons lining the basement membrane following *Arhgap11a* knockdown. (Scrambled: n=3 brains, 1 experiment, *Arhgap11a:* n=4 brains, 1 experiment, unpaired t-test). siRNAs: small interfering RNAs, IUE: *in utero* electroporation, MZ: marginal zone, EM: electronic microscopy, *: p-value<0.05. ***: p-value<0.001 In graphs, individual data points represent different brains. Scale bars: C: 5μm, M, O: 25μm. Bar plots indicate means +/-SEM.

We next evaluated whether knockdown of *Arhgap11a* affected the area of the basal endfoot contacting the basement membrane. Using the 3D reconstructions described above, we observed an average basement membrane coverage of ∼15μm^2^ by control RGC endfeet (**Figure 4F**). However, this coverage significantly diminished by 31% in RGC endfeet depleted of *Arhgap11a*. Of note, there was no observable impact of *Arhgap11a* knockdown on the number of endfeet per RGC (**Figure S4D**). Altogether, these results show that *Arhgap11a* regulates RGC morphology of the basal process and endfeet in the MZ.

This raises the question of what is the impact of these defects upon cortical architecture at the pia? To begin to probe the importance of RGC morphology in the MZ, we used serial-blockface electron microscopy (SBFEM) to visualize RGC basal processes and endfeet structure as well as their interactions with the surrounding MZ niche (**Figure 4G-L**). 3D reconstructions of RGCs together with MZ cells highlighted that basal endfeet tile the basement membrane, forming a tight interface between the MZ and the basement membrane (**Figure 4I**). We also observed RGC basal processes and endfeet in direct contact with the surrounding cells and neurites. For example, we reconstructed tangentially directed cells (presumably interneurons or Cajal Retzius neurons) finding their way through a multitude of RGC basal processes and branches (**Figure 4K,L**). These cells displayed significant contact with basal endfeet tiling the basement membrane.

Given that *Arhgap11a* controls RGC morphology in the MZ, we thus asked whether depleting *Arhgap11a* in RGCs could have a non-cell autonomous effect on these residents of the MZ, including Cajal Retzius neurons and migrating interneurons (Lavdas et al., 1999; Marín and Rubenstein, 2001; Miyoshi and Fishell, 2011; Tanaka et al., 2009; Yokota et al., 2007). We transfected scrambled or *Arhgap11a* siRNAs into RGCs using IUE at E15.5, and analyzed cell density in the MZ 24h later, focusing on the region containing EGFP+ RGC basal processes and endfeet (**Figures 4A,B,M**). First, we used Hoechst staining to quantify MZ cell density. These analyses revealed that compared to control condition, *Arhgap11a* knockdown in RGCs significantly reduced the number of cells lining the LAMININ+ basement membrane (**Figure 4M,N**).

We then quantified interneurons, which migrate tangentially from the ganglionic eminences into the cortex via 3 streams, including along the MZ. Interneurons which migrate through the MZ will invade the CP, along the way forming axons that extend from layer 1 (Lim et al., 2018). This population of interneurons is thus critical for cortical circuitry. To quantify interneurons within the MZ, we employed IUE of siRNAs and EGFP in *Dlx-Cre; Ai14* animals, in which interneurons express the fluorescent protein tdTomato (**Figure 4O**). Strikingly, *Arhgap11a* knockdown in RGCs led to a significant decrease in the number of interneurons lining the basement membrane (**Figure 4O,P**). In contrast, there was no defect in CALRETININ+ cells, which mainly label Cajal Retzius neurons (Tanaka et al., 2009), revealing an exquisitely specific effect of *Arhgap11a* (**Figure S4E-G**). These results demonstrate that *Arhgap11a* acts non-cell autonomously within RGCs to promote the position of a key population of interneurons along the basement membrane. Taken together with the basal process and excitatory neuron phenotypes in the IZ, these findings broadly link RGC morphology in basal structures to non-cell autonomous organization of neurons.

### *Arhgap11a* mRNA is actively transported to radial glial basal endfeet via a 5’ UTR cis element

Our data indicate that *Arhgap11a* is essential for RGC morphology and neuronal migration in two different contexts. Thus, we next sought to interrogate the molecular mechanisms by which *Arhgap11a* controls these RGC sub-cellular compartments, and specifically basal endfeet architecture. Notably, both *Arhgap11a* mRNA and protein are subcellularly enriched in RGC endfeet (**Figure 1**). This suggests that RNA transport and local translation of *Arhgap11a* may enable rapid and specific control of RGC basal endfoot morphology. Active mRNA transport depends on cis localization elements within mRNAs (Huttelmaier et al., 2005). Therefore, we first sought to determine the sequence(s) within *Arhgap11a* mRNA that confers endfoot localization.

To this end, we generated three reporters containing EGFP with a nuclear localization sequence (NLS-EGFP), together with either (i) the 5’UTR (527 bp), (ii) the coding sequence (2,942 bp) or (iii) the 3’UTR of *Arhgap11a* (1,057 bp). To test for position-independent localization capacity, these elements were included in the 3’ end of the reporter, followed by a PolyA sequence (**Figure 5A,B**). The reporters were introduced by *in utero* electroporation at E13.5 and brains were collected one day later at E14.5. As previously shown (Tsunekawa et al., 2012) reporters which are not transported are evident as nuclear EGFP, whereas EGFP localization in the basal process and endfeet indicates that a reporter can be transported from the RGC cell body, and translated in basal endfeet. Using this strategy, we discovered that the endfoot RNA localization sequence of murine *Arhgap11a* mRNA resides in its 5’UTR and not in its coding or 3’UTR sequences (**Figure 5A-C**). This was confirmed by *in situ* hybridization targeting the EGFP sequence in electroporated brains (**Figure S5A,B**).

**Figure 5.**
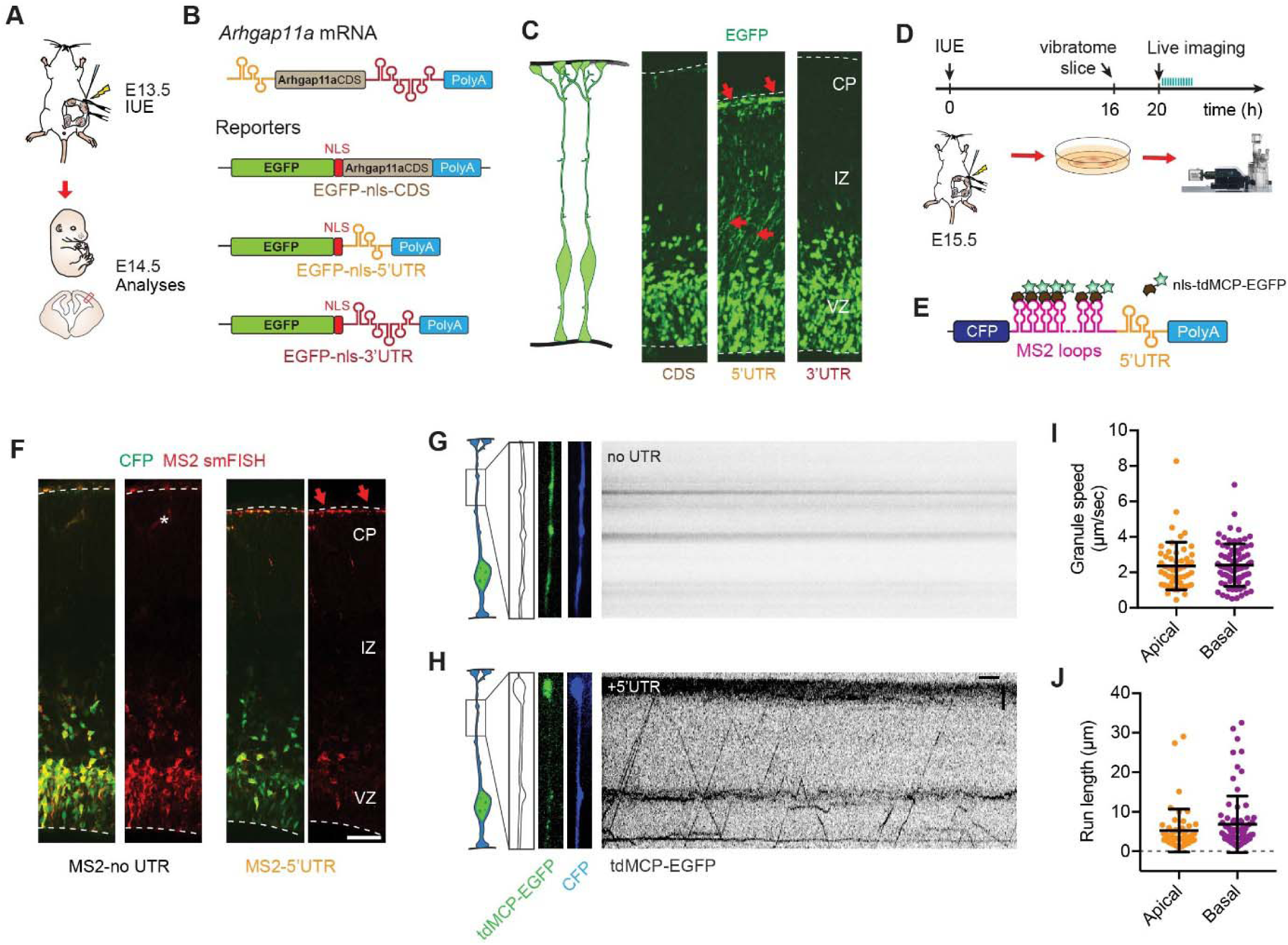
*Arhgap11a* mRNA is actively transported to radial glial basal endfeet via a 5’ UTR element. (A,B) Schematic overview (A) of the strategy used in (B, C) to determine the endfoot localization element in *Arhgap11a* mRNA using a 1-day electroporation of indicated reporter constructs (B). (C) EGFP-nls localizes to RGC basal endfeet only when the *Arhgap11a* 5’UTR is present, but not in CDS alone or containing 3’UTR, thus showing that *Arhgap11a* mRNA endfoot localization sequence resides in its 5’UTR. (D, E) Schematic overview (D) of the strategy used in (F-J) to visualize transport of *Arhgap11a* mRNA reporters (E) in RGC basal processes, using EGFP-labeling of exogenous mRNAs. (F) smFISH (red) targeting MS2 stem-loop RNA sequences shows that addition of *Arhgap11a* 5’UTR induces RNA localization to shift localization from cell bodies to the RGC basal process and endfeet (CFP, green). (G,H) Kymographs showing absence (G) and presence (H) of MS2-tagged mRNA transport in RGC basal process over a 1-min period, in control (noUTR) and 5’UTR, respectively. mRNA is actively transported in RGC basal processes when the tagged mRNA comprises the 5’UTR of *Arhgap11a*. (I,J) Quantification of MS2-tagged mRNA transport in RGC basal processes depicting no average speeds of around 2.5 um/sec in both apical and basally directed movements (I) and similar average run lengths in both types of movements (J). (n=126 EGFP+ punctae, from 11 cells, 2 brains, 2 independent experiments). IUE: *in utero* electroporation, CDS: coding sequence, UTR: untranslated region, nls: nuclear localization signal, tdMCP: tandem MS2-coat protein, CFP: cyan fluorescent protein, CP: cortical plate, IZ: intermediate zone, VZ: ventricular zone. Scale bars: F: 50μm, G,H: horizontal axis: 5sec, vertical axis: 5μm.

Having determined the sequence element of *Arhgap11a* which is sufficient for its localization to endfeet, we next determined if *Arhgap11a* is actively transported to endfeet. To visualize *Arhgap11a* mRNA transport, we performed live-imaging of mRNAs in RGC basal processes within organotypic brain slices (Pilaz et al., 2016a). This technique relies on reporter plasmids containing a CFP expression cassette with a 3’UTR containing 24 MS2 stem loops as well as expression of the nuclear localized tdMCP-EGFP protein with high affinity to RNA MS2 stem loops (**Figure 5D,E**)(Lionnet et al., 2011). By co-transfecting both plasmids this strategy allows the indirect visualization of mRNAs. Two different reporters encoding CFP were generated: (1) one control sequence with MS2 loops only (MS2-no UTR) and (2) the *Arhgap11a* 5’UTR sequence downstream of the MS2 loops (MS2-5’UTR). These reporters were introduced into embryonic brains to label RGCs by *in utero* electroporation at E15.5 and organotypic slices were generated one day later at E16.5. smFISH targeting MS2 loops in fixed electroporated brain sections corroborated localization of the 5’UTR reporter to basal endfeet (**Figure 5F**). This result further confirms that *Arhgap11a* mRNA localization in RGC basal endfeet relies upon its 5’UTR.

Having established these tools, we then performed live imaging of electroporated EGFP+ mRNAs in the basal process of E16.5 RGCs within organotypic brain slices. This allowed us to visualize active directed transport of MS2-5’UTR EGFP+ mRNAs (**Figure 5D,E,G,H, Supplementary Movie 2**). Kymograph analyses highlighted bidirectional movements, with frequent changes of directionality by the 5’UTR reporter but not the no UTR (**Figure 5G,H**). 60% of the observed movements were directed towards the basal endfeet. We measured average speeds of 2.5 μm/s and tracked single movements of up to 32μm, with an average length of 6 μm (**Figure 5I,J**). These speeds and processivity are consistent with active microtubule-dependent transport, and also similar to those observed previously for other mRNA reporters (Pilaz et al., 2016a). Altogether, these results demonstrate that *Arhgap11a* mRNA localization in RGC basal endfeet relies upon active transport and not diffusion along the basal process. This further suggests a specific function for localized *Arhgap11a* in basal endfeet.

### ARHGAP11A protein localization in RGC basal processes and endfeet relies on local translation of *Arhgap11a* mRNA in the basal endfeet

The presence of ARHGAP11A protein in the basal process and endfeet could be explained by at least two possibilities: synthesis of ARHGAP11A in the soma followed by protein transport into the basal process and endfeet, and/or local production of ARHGAP11A within endfeet. Given the active transport of *Arhgap11a* mRNA along the basal process, we predicted that ARHGAP11A protein expression in RGC basal structures relies upon its mRNA localization to endfeet. To test this directly, we generated reporters expressing ARHGAP11A full-length protein fused to the fluorescent photoconvertible protein DENDRA2. Two reporters were generated: (i) a control producing an mRNA devoid of any localization sequence (Dendra2-no UTR) and (ii) another including the 5’UTR of *Arhgap11a* sufficient for RNA localization at the endfoot (Dendra2-5’UTR, **Figure 6A**). DENDRA2 fluorescent signal was analyzed in fixed brains one day after *in utero* electroporation at E15.5. Of note, smFISH targeting *Dendra* confirmed the transport of reporters to basal endfeet (**Figure S6A-C**). Expression of either the control or 5’UTR reporter resulted in DENDRA2 signal in RGC nuclei (**Figure 6B,C**). This cell body localization recapitulates the endogenous protein expression pattern (compare to **Figures 1C-E**). However, DENDRA2 signal in RGC basal endfeet was also evident only in brains electroporated with Dendra2-*Arhgap11a*-5’UTR (**Figure 6B-E**). Further, ARHGAP11A localization was evident along the basal processes within the IZ and CP (**Figure 6D-E**). Localization was not context-dependent as placement of the 5′UTR in either the 5′ or 3′ part of the reporter led to similar enriched localization to basal endfeet **(Figure S6D)**. This strongly argues that ARHGAP11A protein localization both within endfeet, as well as along the basal process, relies upon subcellular localization of the transcript.

**Figure 6.**
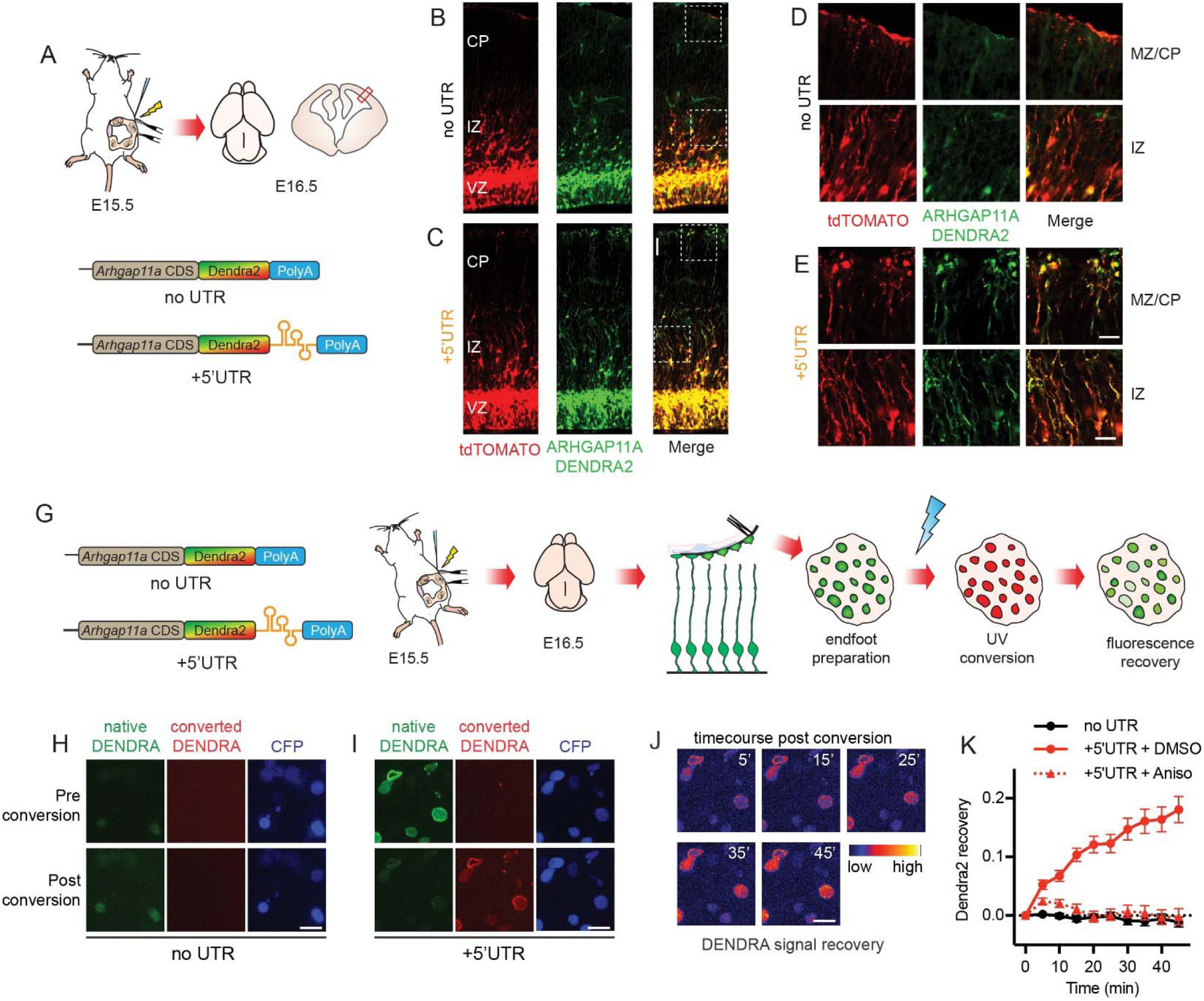
ARHGAP11A protein localization to RGC basal processes and endfeet relies on local translation of *Arhgap11a* mRNA in basal endfeet. (A) Schematic overview of the strategy used in (B-E) to test if *Arhgap11a* mRNA localization mediates ARHGAP11A expression in RGC basal endfeet and basal processes. (B-E) Immunofluorescence of tdTomato electroporated RGCs (red) and ARHGAP11A fusion reporter (green) containing no UTR (top) or 5’ UTR (bottom). High magnification images reflect ARHGAP11A protein localizes to RGC basal endfeet (MZ/CP) and basal processes (IZ) only when *Arhgap11a* 5’UTR is present. (G) Schematic overview of the strategy used in (H-K) to visualize local translation of *Arhgap11a* in RGC basal endfeet. (H-I) Images showing ARHGAP11A-DENDRA fluorescence in RGC basal endfeet pre (top) and post-(bottom) photoconversion in no UTR (H) or 5’ UTR (I) conditions. (J) Time course showing recovery of native DENDRA signal recovery in the +5’UTR condition, as psuedocolored using indicated scale. (time in min). (K) Quantification of positive recovery of native DENDRA signal in RGC basal endfeet only in the 5’UTR+DMSO condition (red, solid line) relative to the no UTR condition (black). Addition of the translation inhibitor anisomycin (red, dotted line) abrogates native signal recovery in the +5’UTR condition. (no UTR: n=27 endfeet from 2 brains, 2 independent experiments, 5’UTR + DMSO: n= 62 endfeet from 3 brains, 3 independent experiments, n=69 endfeet from 3 brains, 3 independent experiments, two-way ANOVA interaction time x condition: p value < 0.0001). UTR: untranslated region, CDS: coding sequence, Aniso: anisomycin. Scale bars: B,C: 100μm; D,E: 20μm; H-J: 5μm. Graph shows average values +/-SEM.

Next, we tested directly if ARHGAP11A can be locally synthesized within endfeet. Using the DENDRA2 constructs described above, together with CFP to identify electroporated endfeet, we employed live imaging to visualize local translation of exogenous *Arhgap11a* mRNAs in CFP+ basal endfeet. Live imaging was performed in *ex vivo* “endfoot preparations” (Pilaz et al., 2016a; Pilaz and Silver, 2017) consisting of isolated basal endfeet connected to the basement membrane and overlying meninges, as previously described (**Figure 6G**). UV exposure was used to photoconvert DENDRA2 signal in isolated endfeet from green to red (**Figure 6H, I**). Subsequently, the recovery of the green fluorescence was monitored over 45 minutes. Since basal endfeet were completely disconnected from the cell body, recovery of any green fluorescence is a robust readout of *de novo* local protein synthesis. No recovery was noted in the negative control lacking the 5′UTR (**Figure 6K**). In contrast, recovery of green fluorescence was observed in endfeet expressing Dendra2-5′UTR over a 45 minute interval (**Figure 6J,K**). Importantly, this recovery was completely abolished when the translation inhibitor anisomycin was added to the culture medium (**Figure 6K**). Altogether, these data demonstrate that *Arhgap11a* mRNA transport and local translation in endfeet are necessary for the sub-cellular localization of the ARHGAP11A protein in RGC basal structures.

### Locally-synthesized ARHGAP11A controls basal process morphology through GAP activity

Endogenous *Arhgap11a* transcript is localized to endfeet beginning around E13.5 and peaking by E15.5, coincident with increasing endfoot morphology. Given the striking localization and timing of both *Arhgap11a* mRNA and protein in basal endfeet, we postulated that local synthesis of ARHGAP11A mediates its functions in endfoot morphology. To test this, we used *in utero* electroporation at E15.5 to introduce *Arhgap11a* siRNAs together with *Arhgap11a* full length protein either containing noUTR or 5’UTR (**Figure 7A-D**). The *Arhgap11a* 5’UTR sequence is sufficient for subcellular localization and translation of ARHGAP11A in the basal process and endfeet (**Figures 6 and S6**). Importantly, both the *Arhgap11a*-noUTR and *Arhgap11a*+5’UTR reporters were resistant to the transfected siRNA cocktail, as the siRNAs specifically targeted *Arhgap11a* 3’UTR. Compared to siRNA knockdown alone, co-expression of *Arhgap11a*-noUTR did not restore the complexity of RGC basal processes at the level of the MZ (**Figure 7E-G,J,K**). In contrast, addition of *Arhgap11a*+5’UTR fully restored the complexity of transfected RGC basal processes in this region (**Figure 7E,G,H,J,K**). Of note, expression of *Arhgap11a*-noUTR or *Arhgap11a*+5’UTR alone (without siRNA) did not affect RGC branching complexity in the MZ, indicating that there is no striking overexpression phenotype (**Figure S7**). These findings demonstrate that subcellular endfoot localization of *Arhgap11a* is critical for proper basal process morphology. Is RGC morphology also dependent upon ARHGAP11A RhoGAP function? To address this question, we generated a GAP-deficient (GD) form of ARHGAP11A, previously described (Zanin et al., 2013). This GD-*Arhgap11a*+5’UTR reporter, contains a single missense mutation, R87A (**Figure 7D**). Similar to the no UTR construct, it also failed to rescue the siRNA-mediated basal process phenotype in the MZ (**Figure 7E,F,I-K**). This demonstrates that local Rho GAP activity, mediated by local synthesis, is essential for basal process complexity in the cortex.

**Figure 7.**
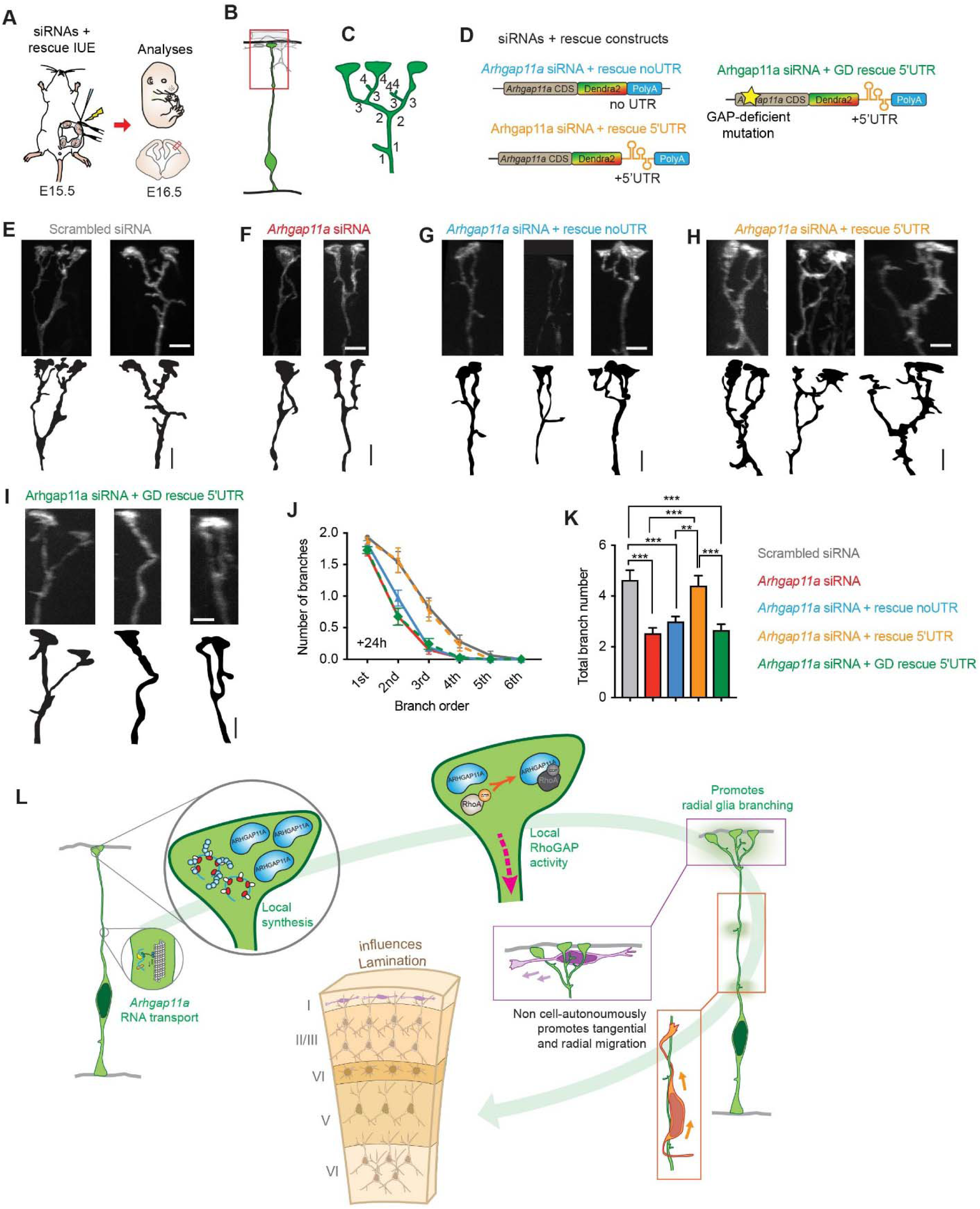
Locally-synthesized ARHGAP11A controls basal process morphology through GAP activity. (A) Schematic overview of the strategy used in (B-K) to assess rescue of RGC endfeet morphology. (B,C) Method used to define branch orders in in RGC basal processes in MZ. (D) Rescue constructs used in experiments. (E-I) Representative images showing basal process complexity at the level of the MZ in RGC treated with different conditions. (J,K) Quantification of basal process and endfoot complexity at the level of the MZ in RGC treated with different conditions. (Scrambled: n=78 cells from 6 brains, 4 independent experiments, *Arhgap11a:* n=78 cells from 5 brains, 4 independent experiments, *Arhgap11a* + rescue: n= 97 cells from 6 brains, 4 independent experiments, *Arhgap11a + rescue 5’UTR*: n= 56 cells from 4 brains, 3 independent experiments, *Arhgap11a* + GD rescue 5’UTR: n=78 cells, from 4 brains, 2 independent experiments, J: 2-way ANOVA: p-value<0.0001, K:One way ANOVA: p<0.0001, Tukey’s Post-Hoc comparisons). (L) Cartoon representation of the major findings of this study. *Arhgap11a* mRNA is actively transported in RGC basal process to basal endfeet. In basal endfeet local synthesis of ARHGAP11A protein enables expression in the basal process and local RhoGAP activity, thus promoting radial glia branching. Thus, mRNA localization and local translation (of Arhgap11a) have a non-cell autonomous role in promoting migration of both excitatory and inhibitory neurons, which ultimately impacts post-natal cortex lamination. IUE: *in utero* electroporation, siRNA: small interfering RNA, CDS: coding sequence, UTR: untranslated region, GD: Rho-gap-deficient. **: p-value<0.01. ***: p-value<0.001. E-G: 5μm. Graphs show average values +/-SEM.

Altogether, these results demonstrate that (1) *Arhgap11a* employs mRNA transport and local protein synthesis at RGC endfeet to promote complex basal process morphology in the MZ, (2) this regulation relies on its GAP activity and suggest that (3) *Arhgap11a*-dependent control of RGC morphology regulates neuron positioning in the cerebral cortex (**Figure 7L**). This establishes an essential and novel function for subcellular gene expression in radial glia and the regulation of cortical development.

## Discussion

By controlling neurogenesis and radial migration, RGCs plays a crucial role in orchestrating the development of the cerebral cortex. Yet how the polarized morphology of RGCs is precisely controlled to dictate these functions is poorly understood. Here, we demonstrate that RGC morphology and thus cortical architecture relies upon exquisite temporal and spatial control of gene expression via active transport and local translation of the RhoA-GAP *Arhgap11a*. This subcellular gene expression enables *Arhgap11a* to function non-cell autonomously in RGCs to direct tangential migration of interneurons in the MZ, and radial migration of excitatory neurons in the IZ. Thus, RhoA-GAP activity is spatially and acutely activated via local translation in RGCs to mediate neuron positioning and cortical cytoarchitecture. This demonstrates, for the first time, that mRNA localization and local translation in RGCs controls brain development *in vivo*. Thus, our study implicates a dynamic new gene regulatory mechanism by which progenitors shape organismal development and highlights a valuable *in vivo* model for understanding how local gene regulation controls complex cellular functions.

### mRNA transport and local translation are essential for RGC morphology

Using live and fixed imaging both *ex vivo* and *in vivo*, we demonstrate that *Arhgap11a* undergoes active transport, subcellular localization and local translation in RGC basal endfeet. This discovery adds *Arhgap11a* to a short but growing list of transcripts which we and others have shown undergo subcellular localization and trafficking in RGC basal processes (Dahlstrand et al., 1995; Pilaz et al., 2016a; Tsunekawa et al., 2012; Tsunekawa et al., 2014). This further reinforces the discovery that RGC basal processes are highways for active mRNA transport (Pilaz et al., 2016b; Pilaz and Silver, 2017).

Most importantly, we demonstrate that at least one function of subcellular RNA localization in RGCs is to mediate local morphology. This is based upon several pieces of evidence. First, we observed that exogenous ARHGAP11A RNA and protein localization to basal endfeet and basal processes relies entirely upon its 5’UTR, and coincides with critical developmental stages when RGC branching becomes more complex (Lu et al., 2015). Second, acute depletion of *Arhgap11a* decreases RGC branching complexity in the basal process and endfeet; importantly and strikingly, this phenotype was rescued only by concomitant expression of endfoot-localized *Arhgap11a* mRNA. Third, we show that RGC branching depends upon localized RhoGAP activity. These data clearly demonstrate that RGC basal structures rely upon active mRNA transport, local translation, and local Rho GTPase signaling.

Why do RGCs utilize new protein production in endfeet rather than simply transport proteins from the cell body? In polarized neurons and migratory cells, RNA transport and local protein synthesis mediate rapid and sub-cellular functions (Buxbaum et al., 2015). Translation of an average protein takes about 1 minute, whereas transcription persists over 10 minutes (Shamir et al., 2016). Thus, local expression is significantly more efficient and energetically favorable, enabling rapid changes in RGC morphology as the brain grows radially and tangentially. Beyond *Arhgap11a*, the endfoot transcriptome contains many mRNAs encoding cytoskeletal and signaling regulators (Pilaz et al., 2016a), suggesting this may be a widespread mechanism in RGCs. Of note, migrating fibroblasts and neuronal growth cones and spines also contain similar classes of localized transcripts (Cajigas et al., 2012; Goering et al., 2019; Poulopoulos et al., 2019). Yet, control of RGC morphology may be just one of many functions of mRNA localization in endfeet. Roles for localized RNA in cell fate have also been postulated (Tsunekawa et al., 2012). It will be exciting to interrogate how additional localized transcripts coordinate with *Arhgap11a* to control RGC complexity as well as influence other dynamic events in the basal process and endfeet, such as signaling with the surrounding niche.

### Localized Rho-GTPase signaling controls RGC morphology

Our findings demonstrate that exquisite spatial and temporal control of Rho GTPase signaling is instrumental for RGC morphology. Rho signaling has been previously linked to RGC morphology, but how this was controlled rapidly and spatially was unknown. Global manipulation of GTPases disrupts filopodial activity in RGC basal structures, ultimately impairing neuronal migration and neurogenesis (Haubst et al., 2006; Li et al., 2008; Radakovits et al., 2009; Yokota et al., 2010) Further, RhoA cKO or disruption of a Rho modulator, PLEKHG6, impairs RGC organization and non-cell autonomously disrupts neuronal migration (Cappello et al., 2012; O’Neill et al., 2018). In neurons, RhoA inactivation increases branching, whereas RhoA activation reduces branching (Lee et al., 2000). Our findings suggest parallel mechanisms are at play in RGCs, where RhoA overactivation via ARHGAP11A loss also decreases branching. Given its characterized roles, ARHGAP11A may ultimately influence both actin and microtubule networks (Müller et al., 2018; Zanin et al., 2013). Our findings suggest broadly that localized pools and expression of GTPase regulators like ARHGAP11A enable exquisite subcellular control of ubiquitous GTPases.

How then does locally synthesized endfeet ARHGAP11A influence complexity along the basal process? We posit that ARHGAP11A which is translated in endfeet moves into the basal process either passively or actively. In support of this, ARHGAP11A protein and reporters localized to the basal process both in the MZ as well as the IZ/CP, but only when the endfoot localization element was present (**Figures 1, 6**). Notably, we also observed sporadic enrichment of *Arhgap11a* mRNAs along the basal process (**Figures 1I, 5F**). While this could be a result of collective RNA trafficking to basal endfeet, such enrichment could also represent local translation “hotspots” in the basal process, as proposed in axons (Kim and Jung, 2020).

### Link between localized *Arhgap11a* expression, basal process morphology, and migration

*Arhgap11a* expression in RGCs non-cell-autonomously promotes neuronal migration. Loss of *Arhgap11a* led to more static neurons, fewer neurons migrating basally and an overall reduction in net distance travelled. Importantly, we showed this was not due a requirement of *Arhgap11a* for cell fate nor its expression within neurons. This non-cell-autonomous effect is rapid and long-lasting, given phenotypes manifest following an acute one-day depletion of *Arhgap11a*, yet the lamination defects are evident 25 days later. Given roles for *Arhgap11a* in basal process and endfoot morphology, we propose that these defects cause impaired neuronal migration. It is possible that ARHGAP11A also controls neuronal migration by affecting extracellular signaling and/or cell surface molecules. However, importantly, our observations are in line with studies of RhoA signaling which also modulates RGC morphology and non-cell-autonomously impairs neuronal migration (Cappello et al., 2012). Likewise, knockdown of *Memo1*, another factor linked to RhoA signaling (Zaoui et al., 2008), causes uneven distribution (tiling) of RGC basal processes, hyperbranching, and migration defects (Nakagawa et al., 2019). Altogether this indicates that both simplified and excessive RGC branching may impair neuronal migration and disrupt lamination.

Of note, our study is the first to demonstrate a link between RGC morphology and interneuron migration. Interneurons traverse the developing cortex mainly via the VZ and MZ and the ultimate destination of interneurons is tied to their fate (Bartolini et al., 2017). The population of migrating interneurons in the MZ are specialized, developing into Martinotti cells which also leave axons in layer I as they invade the CP (Lim et al., 2018). Interneuron mis-positioning in the MZ was evident with just a one-day acute depletion of *Arhgap11a* from RGCs. It will be interesting to test whether more sustained manipulation of *Arhgap11a* localization in RGCs and thus, disruption of RGC morphology, will impact cortical circuitry.

### Implications for human cortical development and disease

During evolution, a hominid-specific partial duplication of *ARHGAP11A* led to the emergence of *ARHGAP11B*, which has been linked to cerebral cortex expansion in humans. In the human developing cortex, *ARHGAP11B* is expressed in neural progenitors and its forced-expression in developing mouse cortices, ferrets and marmosets promotes basal progenitor generation and neuron production (Florio et al., 2015; Heide et al., 2020; Kalebic et al., 2018). *ARHGAP11B*, however, lacks both the GTPase activity and the RNA motif for localizing to endfeet and indeed shows no subcellular localization in human cortices (Florio et al., 2018). It is interesting to consider the relationship between *ARHGAP11A* and *ARHGAP11B* in human RGCs; whereby the human-specific form could modulate the ancestral protein’s function in the cell body, similar to that seen for other human-specific duplications like SRGAP2 (Charrier et al., 2012) or NOTCH2NL (Fiddes et al., 2018; Suzuki et al., 2018).

Given the conserved localization of *ARHGAP11A* in the human neocortex, we predict that this RhoGAP and more broadly, mRNA localization, is critical in human RGCs. In human neocortices, the RGC basal process is significantly longer than in mice (up to several millimeters) and a prominent feature of outer radial glia (oRGs), a major progenitor in primates (Betizeau et al., 2013; Florio et al., 2015; Hansen et al., 2010). Indeed, *ARHGAP11A* is highly expressed in both human RGCs and oRGs. Thus, localized pools of *ARHGAP11A* mRNA and protein may promote human cortical development by influencing RGC and oRG morphology. Indeed, in primates, lamellar expansions decorate the basal process, and reside in close contact with migrating neurons (Rakic, 1972). It is also important to note that defects in RGC scaffolds underlie diverse neurodevelopmental diseases including heterotopias and lissencephaly (Lambert de Rouvroit and Goffinet, 2001; Myshrall et al., 2012). This highlights the importance of investigating subcellular mRNA localization and local translation in RGCs, towards understanding both normal development and disease.

## Acknowledgements

The authors thank members of the Silver lab and Denis Jabaudon for discussions and careful reading of the manuscript. This work was supported by NS083897, NS099350, MH119813 (to DLS). We also thank Duke EM and imaging core facilities.

## Experimental Procedures

### Mice

All experiments were performed in agreement with the guidelines from the Division of Laboratory Animal Resources from Duke University School of Medicine and approved by Duke IACUC. Plug dates were defined as embryonic day (E) 0.5 on the morning the plug was identified. All experiments were conducted in the C57BL/6J strain. The following mouse strains were used: Dlx-Cre (Tg(dlx5a-cre)1Mekk/J); Ai14-tdTomato (B6.Cg-Gt(ROSA)26Sor^tm14(CAG-tdTomatoHze^/J) both from Jax labs).

### Human embryonic samples and *in situ* hybridization

The study using human fetal sample (9 wpc) was approved by three relevant Ethics Committees (Erasme Hospital, Université Libre de Bruxelles, and Belgian National Fund for Scientific Research FRS/FNRS) on research involving human subjects. Written informed consent was given by the parents in each case. *In situ* hybridization (ISH) on human fetal cortical sections using digoxigenin-labeled RNA probes was performed as described previously (Suzuki et al., 2018). The probe specifically recognizing exon11 and 12 of human ARHGAP11A, not ARHGAP11B, was prepared (2117– 2657 of human ARHGAP11A (NM_014783.5)). Imaging was performed using a Zeiss Axioplan 2 and the intensity and contrast were modified using Fiji/ImageJ software. We confirmed the specificity of the signal produced by the anti-sense pan-ARHGAP11A probe by comparing with the virtual absence of the signal by the sense probe.

### Histology

Mouse embryos were collected and dissected in cold phosphate buffer saline (PBS) and histology was performed as previously (Pilaz et al., 2016a). Brain fixation was performed overnight by immersion in a 4% Paraformaldehyde 1X PBS solution. Following fixation, embryonic brains were washed twice in cold PBS for 20min. Cryoprotection was performed by overnight immersion in a 30% sucrose (w/v) PBS solution. Following cryoprotection, brain were transferred into OCT medium. Cryosections were generated using a cryostat and deposited on glass slides. The thickness of the sections varied depending on the purpose of the experiments (10-20μm for characterization of ARHGAP11A localization by immunofluorescence and *in situ* hybridization, 50μm for characterization of basal process morphology with 3D reconstructions, 30μm for analysis of cortical layering at post-natal stages).

Immunofluorescence was performed as described previously (Pilaz et al., 2016a). Briefly, slides were left to thaw at room temperature (RT) for 10min. Then, they were washed by immersion in PBS for 10min, followed by permeablization with a 15-30min wash in 0.2-0.5% Triton-X (w/v) in PBS. Following this step, sections were washed once in PBS and blocking was performed with Mouse on Mouse (MOM, Vector Laboratories) blocking agent when using primary antibodies produced in the mouse, and 10% NGS for primary antibodies generated in any other species. Primary antibody incubation in PBS or MOM diluent was then performed overnight at 4C. Three 5-10-minute washes were then performed in PBS, followed by 30-60min RT incubation in a secondary antibody solution containing Hoechst. Before mounting was performed in Vectashield (Vector Laboratories) or Mowiol, three 5-10-minute washes were performed in PBS. Primary antibodies used in this study were the following: rabbit anti-Arhgap11a (Bethyl, A303-097A), rabbit anti-Arhgap11a (Abcam, ab113261), rat anti-Nestin (BD biosciences, 556309), rabbit anti-Tbr1 (Abcam, ab31940), rat anti-Ctip2 (Abcam, ab18465), mouse anti-Ror-Beta (R&D Systems, N7927), rabbit anti-Laminin (Millipore, AB2034), rabbit anti-Calretinin (Swant, 7697), rat anti-Sox2 (Thermo Fisher, 14-9811-80), rabbit anti-Tbr2 (Abcam, ab183991), rabbit anti-Ki67 (Cell signaling, 12202). Cell counting was performed in FIJI (ImageJ), using the *Cell counter* plugin.

For binning analyses, X-Y coordinates were extracted from *Cell counter* data, and a script created in R (R Development Core Team, 2019) was used to assign punctae to specific bins, using the coordinates of the ventricular and pial borders as references. Bin numbers were reported in an Excel spreadsheet for analyses.

### DNA constructs and siRNAs

pCAGGS-EX and pCAGGS-EGFP plasmids were kind gifts from Nicholas Gaiano (Mizutani et al., 2007), the pGLAST-EGFP-CAXX plasmid was kindly offered by Tarik Haydar(Gal et al., 2006), and the pCAGGS-PB-mCherry-CAXX was generously given by Cagla’s Eroglu laboratory. EGFP-nls-CDS, EGFP-ns-5’UTR and EGFP-nls-3’UTR constructs were generated by cloning sequences of interest downstream of the EGFP sequence, using *Arhgap11a* cDNA cloned from mouse embryonic cortical cDNA as described previously(Tsunekawa et al., 2012). The MS2-no UTR plasmid was described previously (Pilaz et al., 2016a). We used a Gibson assembly strategy (NEB Hifi Builder) to clone *Arhgap11a*’s 5’UTR from the EGFP-nls-5’UTR construct into the MS2-no UTR plasmid downstream of the MS2 stem loops sequence. Similarly, we used Gibson assembly to generate the Dendra2-no UTR and the Dendra2-5’ UTR plasmids, cloning 2 fragments (*Arhgap11a* and *Dendra2* coding sequences) or three fragments (*Arhgap11a* and *Dendra2* coding sequences followed by *Arhgap11a 5’UTR*) into the EcoRI site of pCAGGS-EX, using EGFP-nls-CDS, Dendra2-no UTR (Pilaz et al., 2016a), and EGFP-nls-5’UTR as templates, respectively. siRNAs targeting the 3’UTR of *Arhgap11a* were purchased (siRNAflex, Qiagen).

### *In utero* electroporation (IUE)

We performed IUE as described previously (Pilaz et al., 2016a; Saito and Nakatsuji, 2001). Electroporation parameters were as follows: five consecutive 50ms electrical pulses spaced by 950ms, voltage varied from 40V to 60V depending on the embryonic stage at which the procedure was performed. Plasmids were produced using Qiagen or Sigma Endotoxin Free Maxi Prep kits and following the manufacturers’ instructions. Individual plasmid concentrations injected into lateral ventricles ranged from 0.5 to 1μg/ul. siRNAs were injected at a final concentration of 2.5μM.

### Single molecule fluorescent in situ hybridization (smFISH)

*In situ* hybridization was performed following the protocol described by Takahashi and Osumi (Takahashi and Osumi, 2002). Probes sequences are listed in *Supplementary table 1*. The protocol used to reveal *Arhgap11a, MS2* or *Dendra2* mRNAs by smFISH was as previously (Pilaz et al., 2016a). All buffers and solutions used for this protocol were previously treated with diethyl pyrocarbonate to quench RNA-ase activity. A pool of Stellaris probes targeting *Arhgap11a* and labeled with the Quasar570 fluorophore were purchased from Biosearch. *MS2* and *Dendra2* probes were described previously (Pilaz et al., 2016a). For quantification of *Arhgap11a* mRNA density after IUE-mediated siRNA knockdown, 10μm mosaic Z-stacks covering the entire thickness of the electroporated and the corresponding region in the non-electroporated hemisphere were acquired using a 63X objective with a microscope equipped with Apotome technology (Zeiss). The electroporated region was evident based on the presence of EGFP+ cells. Coordinates of smFISH punctae were manually registered using the *Cell Counter* plugging in FIJI (ImageJ, over one thousand punctae were registered in non siRNA treated regions). A script created in R (R Development Core Team, 2019) was used to assign punctae to specific bins. Bin numbers were reported in an Excel spreadsheet.

### qPCR analyses in Dcx-DsRed embryos

Cortices from E14.5 *Dcx*::DsRed embryos were isolated, incubated with 0.25% trypsin-EDTA solution for 10 min. at 37°C, dissociated into a single cell suspension, and sorted in a Sorter Astrios machine. Positive and negative cells were directly collected into RNA extraction buffer (RLT) supplemented with 1% β-Mercaptoethanol. Samples were vortexed and RNA was extracted using Qiagen RNeasy plus kit. cDNA was synthesized from RNA using Biogen iScript kit and qPCR was performed using either Sybr Green iTaq (BioRad) or TaqMan (Life Technologies) in an Applied Biosystems StepOne machine (Thermo Fisher Scientific). The following primers and TaqMan probes were used in the qPCR reaction: β*-Actin* (5’ Forward-AGATCAAGATCATTGCTCCT and 3’ Reverse-CCTGCTTGCTGATCCACATC), *Pax6* (5’ Forward-TCTTTGCTTGGGAAATCCG and 3’ Reverse-CTGCCCGTTCAACATCCTTAG), *Arhgap11a* (5’ Forward-GCAGGTGTGCCAAGGCGAAGT and 3’ Reverse-TGCAAGTCGCCAACCAACACTTTCA)(Kagawa et al., 2013), *Gapdh* (Mm99999915_g1), Tubb3 (Mm00727586_s1). Values were normalized to *Gapdh* (TaqMan) or β*-Actin* (Sybr Green) as loading control.

### Live imaging

We performed and quantified live imaging of RNA trafficking, using methods identical to those described previously (Pilaz et al., 2016a). For imaging of translation, following *in utero* electroporation of *Dendra2* and pCAGGS-CFP plasmids at E15 in the afternoon, brains were dissected in cold 1x HBSS supplemented with 2.5 mM HEPES, 30 mM D-glucose, and 4mM NaHCO3 during the morning on the next day. We generated endfoot preparations as described previously (Pilaz et al., 2016a), using tweezers to peel off the basement membrane together with connected endfeet from the surface of the brain. 2-4 endfoot preparations from brains electroporated with different plasmid conditions were mounted together in a 1mg/ml collagen solution supplemented with DMEM/F12 at the bottom of 35-mm glass bottom dishes (MatTek). A slab of 3% agarose gel was added on top of endfoot preparations to prevent their detachment from the bottom of the dish. Additional collagen was added to stabilize the preparation. Endfoot preparations were cultured in DMEM-F12 supplemented with B27 without vitamin A (Gibco), N2 (Gibco), 5% horse serum, 5% fetal bovine serum, 10ng/ml FGF and 20ng/ml EGF. Culture medium was added after a 15min incubation at 37C to ensure proper polymerization of collagen matrix. Endfoot preparations were left to equilibrate at 37C and 5% CO2 for 1-2h prior to live imaging. Live imaging was performed with a 100x/1.4 oil U PlanSApo objective mounted on an inverted spinning disk confocal microscope (Andor XD revolution spinning disk confocal microscope), equiped with a 37C and 5% CO2 incubation chamber. Following a one-hour equilibration of the incubation chamber, three 15μm Z-stacks per endfoot preparations were imaged in the blue, green and red channels with a 2μm resolution in the Z axis (pre-conversion acquisitions). This allowed the simultaneous recording of several conditions within one session and therefore minimized variability between imaging sessions. Positions were selected at locations where endfoot preparations presented minimal folding. Each position was then exposed to Arc lamp illumination at 10% intensity for 20 seconds with manual scanning in the Z-dimension. Each Z-stack was then acquired in the blue, green and red channels (post conversion acquisitions). Z stacks were then acquired every 5min for 45min in the green and blue channels (timecourse acquisitions). Following this, 40μM anisomycin treatment was performed for 20min and positions unaffected by the intial photoconversion were imaged using parameters described above (Aniso acquisitons). The FIJI software was used for the quantification of green fluorescence signal recovery over time. Z-projections were generated for all the Z-stacks. The ellipse-selection tool was used to generate regions of interests (ROIs) covering individual endfeet at each time point. Average green fluorescence intensity in endfeet was reported at each timepoint into an excel table where Dendra2 recovery was calculated using the following formula: (Dendra2 recovery)_t_ = [(Dendra2 signal)_t_ -(Dendra2 signal)_t0_] / (Dendra2 signal)_t0_.

### Analyses of basal process branching in the MZ

Slides containing sections from brains electroporated with pGLAST-EGFP-CAXX, siRNAs and rescue plasmids were washed once in PBS for 10min, followed by a 15min wash in 0.25% Triton X in PBS (w/v) and one additional PBS wash for 15min. Vectashield was used to mount the slides with a coverslip and prevent bleaching of the EGFP signal. 20-40μm Z-stack images were acquired at a 0.2μm resolution in the Z dimension, using a 63X objective mounted on an epifluorescence microscope equipped with the Apotome technology (Zeiss). Analyses were performed in FIJI. Maximum intensity Z-projections of the Z stacks were generated to identify entire basal processes and endfeet. Rectangular regions of interests (ROIs) were traced around individual or small groups of basal processes. 3D projections of selected ROIs were generated with the following parameters deviating from the default settings: “axis of rotation”: *Y axis*, “rotation angle increment”: *5*, “Interpolate”: *checked*. We used the *Cell Counter* plugin to count branches of each order, rotating the 3D projection in order to identify branches obscured by other basal processes. First order branches were those located further away from the pial surface (see Figure 4 for a depiction of branch orders). We used the *Line* tool to ensure that we counted branches >5μm only. The number of branches of each order was then reported into an Excel spreadsheet and the total number of branches was calculated as the sum of all the branches from different orders.

### Analyses of endfoot area covering the basement membrane

These analyses were performed with Z-stack images used for analyses of basal process branching in upper CP/MZ regions. To quantify endfoot area, Z-stack images were first rotated so that the membrane would be orientated in the X-axis. Using Maximum Intensity Z-projections, rectangular regions of interest comprising a small number of endfeet were traced close to pial border. 3D projections of these regions were generated as described above, with the exception that in this case, the “axis of rotation” was set to *X axis*. 3D projection images were then rotated until endfoot areas were the largest, thus truly reflecting actual area covering the basement membrane. The *Polygon Selection* tool was then used to trace the outside limits of basal endfeet, and measured areas were reported into an Excel spreadsheet.

### Analyses of basal process extensions in the IZ-low CP regions

These analyses were very similar to those performed in the upper CP/MZ regions, with the following modifications. Z-stack images of basal process in the IZ, low CP regions were acquired on a LSM 710 Zeiss confocal microscope. Here we did not assess the order of branches and a 5μm cutoff was not used. Instead we counted the total number of extensions emanating from the acquired region of the basal process and normalized this number to the length of basal process that we acquired and focusing on processes with a visualized length >50μm. Rotation of 3D projections were still used to visualize extensions masked by other structures.

### Live imaging of neuron migration

A first *in utero* electroporation (IUE) of a pCAGGS-EGFP plasmid was performed at E14.5. 24h later, another IUE procedure was performed to transfect siRNAs together with a pCAGGS-PB-mCherryCAXX into the hemispheres transfected earlier with pCAGGS-EGFP. In the morning of the next day, 300μm-thick organotypic brain slices were generated as described previously(Pilaz and Silver, 2014). Slices were cultured on cell culture inserts, and were surrounded with a 1mg/ml collagen solution supplemented with DMEM-F12. We used the same medium as the one described above for Dendra2 live imaging of basal endfeet. 4-6 slices transfected with different siRNA conditions were mounted on the same inserts. In the evening, live imaging was initiated for a total of 16h. Z-stack images of regions showing clear overlap between the first and second electroporations were acquired every 10min, using a 10x objective mounted on an inverted LSM 710 Zeiss confocal microscope, equipped with an incubation chamber (37C, 5% CO2). Analyses were performed in FIJI, using the *Manual Tracking* plugin. We focused on neurons located in the low IZ / SVZ region at t_0h_. Tracking results were reported in an Excel spreadsheet and the analyzed parameters are defined as follows. Net distances in Y and X: |*Y*_*0h*_*-Y*_*16h*_| and |*X*_*0h*_*-X*_*16h*_|, respectively. Net distance: Euclidean distance between neuron positions at t_0h_ and t_16h_. Total path length: sum of distances traveled between each time points. Velocity: average travel speed between each time point. For analysis of movements in Y as “down”, “steady”, or “up”, we calculated the Y distance traveled from one time point to the next (Y_t1_-Y_t2_). “Down”, “steady” and “up” were defined as negative, null and positive Y distances, respectively.

### Serial Block Face Electron Microscopy

For SBF-SEM, the samples then underwent a heavy metal staining protocol adapted from Deerinck (2010) after fixation. The tissues were washed in 0.1M sodium cacodylate pH7.4 and then a solution of 1.6 % potassium ferrocyanide with 2% osmium tetroxide buffered with 0.1M sodium cacodylate was added for 1h at RT. This is followed by filtered 10% thiocarbohydrazide (TCH) freshly prepared for 30min. Samples were then washed in distilled water and a secondary 2% osmium tetroxide incubation for 30min. The samples were then placed in 1% uranyl acetate at 4C overnight, washed in distilled water and then placed in freshly prepared lead aspartate solution for 30min at 60C. The samples were then dehydrated with cold ethanol, from 25% to 100% and then infiltrated with increasing concentrations of Durcupan resin in ethanol with several exchanges of 100% resin. The samples were finally embedded in 100% resin and left to polymerize at 60°C for 48 hours.

The tissue samples embedded in resin were manually trimmed with a razor blade to expose the tissue on their surfaces, and then glued onto an aluminum SBF-SEM rivet with conductive epoxy (SPI Conductive Silver Epoxy) with the exposed tissue down. Specimens on the rivet were further trimmed with a razor blade to as small a size as possible (about 0.5mm), and block face was trimmed with a glass knife. Once tissue was exposed, semi thin sections 0.5μm were taken and stained with toluidine blue and viewed under a light microscope to check tissue orientation and condition. Then, the rivet with the sample was sputter coated with gold-palladium.

The image stacks were acquired in an automated fashion by using a high resolution scanning electron microscope (Merlin - Carl Zeiss, Germany) equipped with a 3View system (Gatan Inc., Pleasanton, CA, USA), and a back-scattered electron detector. The Digital Micrograph software (Gatan Inc.) was used to adjust the SEM imaging conditions and slicing parameters. The electron microscope was operated at the high-resolution mode with an acceleration voltage of 2 kV, current mode and in the high-vacuum mode. All images were taken with the following scanning settings at 80pA, dwell time = 2s; pixel sizes 5-7nm. On average 300 sections were obtained at 60nm thickness

## Statistical analyses

All analyses were performed in a blinded fashion. Number of data points and statistical tests used for all the comparisons are indicated in the figure legends.

---

**Figure S1.**
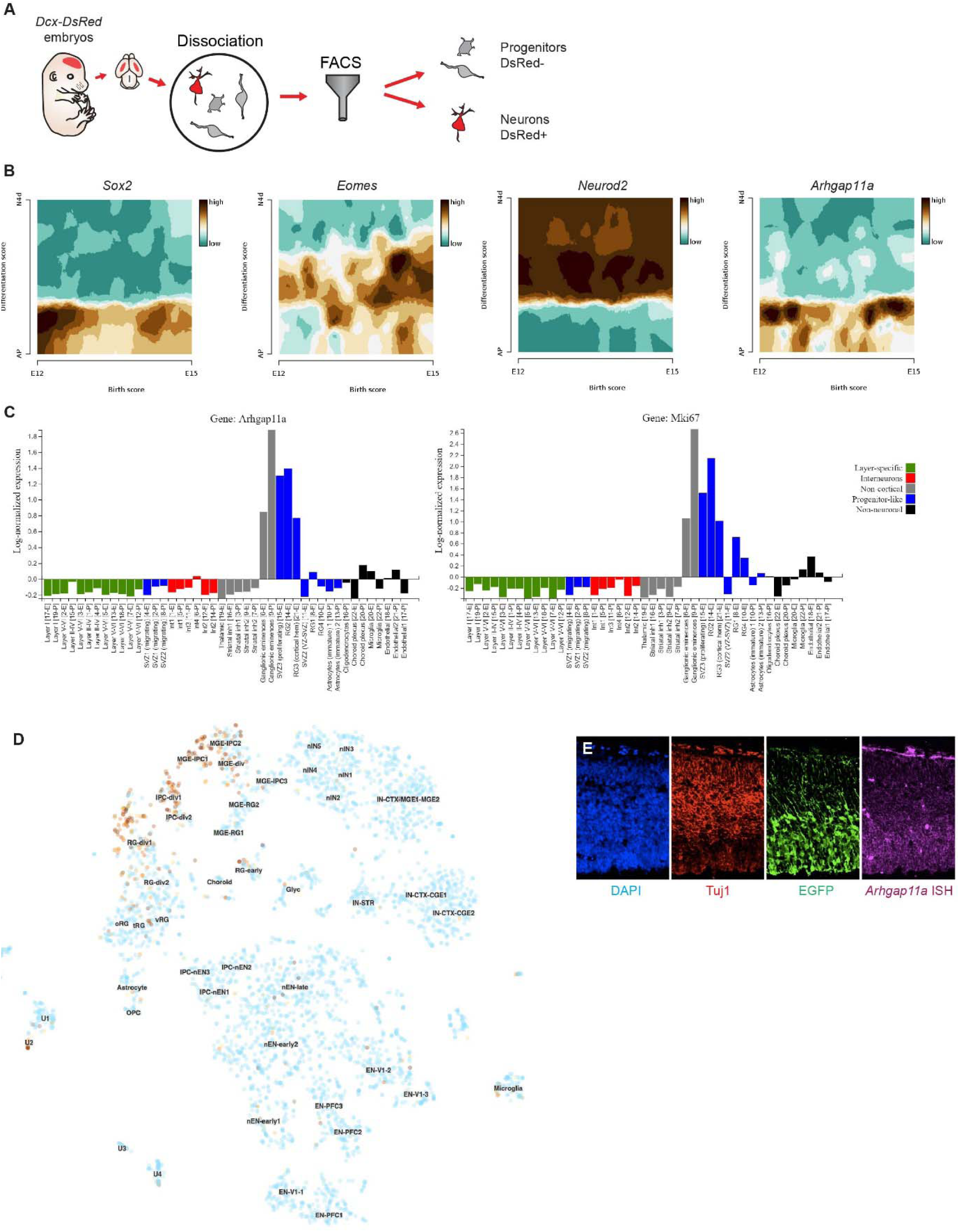
*Arhgap11a* expression is restricted to neural progenitors in mice and humans. Related to Figure 1. (A) Cartoon showing the strategy used to perform qPCR analysis of mRNA levels in sorted embryonic cortical cells in Figure B. (B-D) Published single-cell RNA-sequencing analyses confirm *Arhgap11a* expression is restricted to neural progenitors in mice from E12 to E15 (A): (Telley et al., 2019), in mice at E14.5 and P0 (B): (Loo et al., 2019), and in human fetal cortices from peak neurogenesis stages (C) (Nowakowski et al., 2017). (E) *In situ* hybridization coupled with immunofluorescence in E14.5 EGFP-electroporated brains shows strong enrichment of *Arhgap11a* mRNA (purple) in EGFP+ basal endfeet (green)and not in Tuj1+ neurons (red). FACS: fluorescence-activated cell sorting, ISH: in situ hybridization, EGFP: enhanced green fluorescent protein.

**Figure S2.**
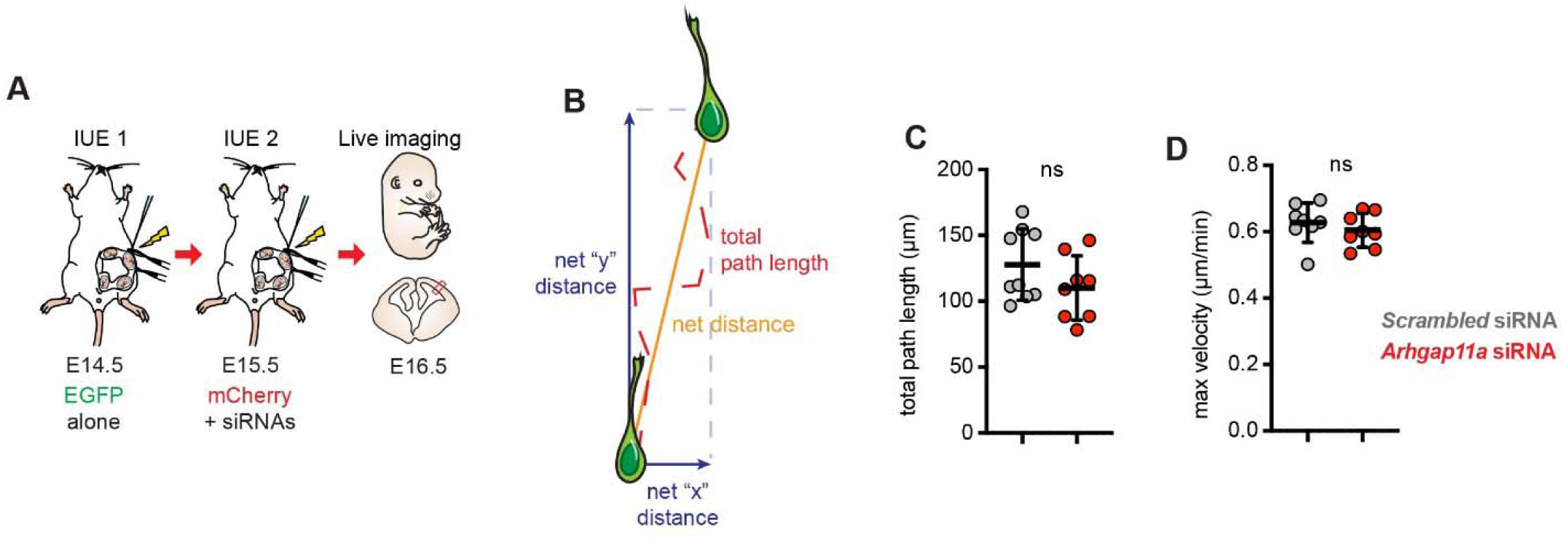
Neuronal migration parameters unaffected by depletion of *Arhgap11a* in RGCs. Related to Figure 2. (A) Schematic overview of experiments to test the impact of *Arhgap11a* depletion in RGCs on neuronal migration. (B) Neuronal migration parameters analyzed. (C,D) Quantification of neuronal migration parameters, reflecting no defect in total path length nor max velocity following depletion of *Arhgap11a* in RGCs. (Scrambled and *Arhgap11a*: n=8 brains from 2 independent experiments, unpaired t-tests) IUE: *in utero* electroporation, siRNA: small interfering RNA, ns: non-significant.

**Figure S3.**
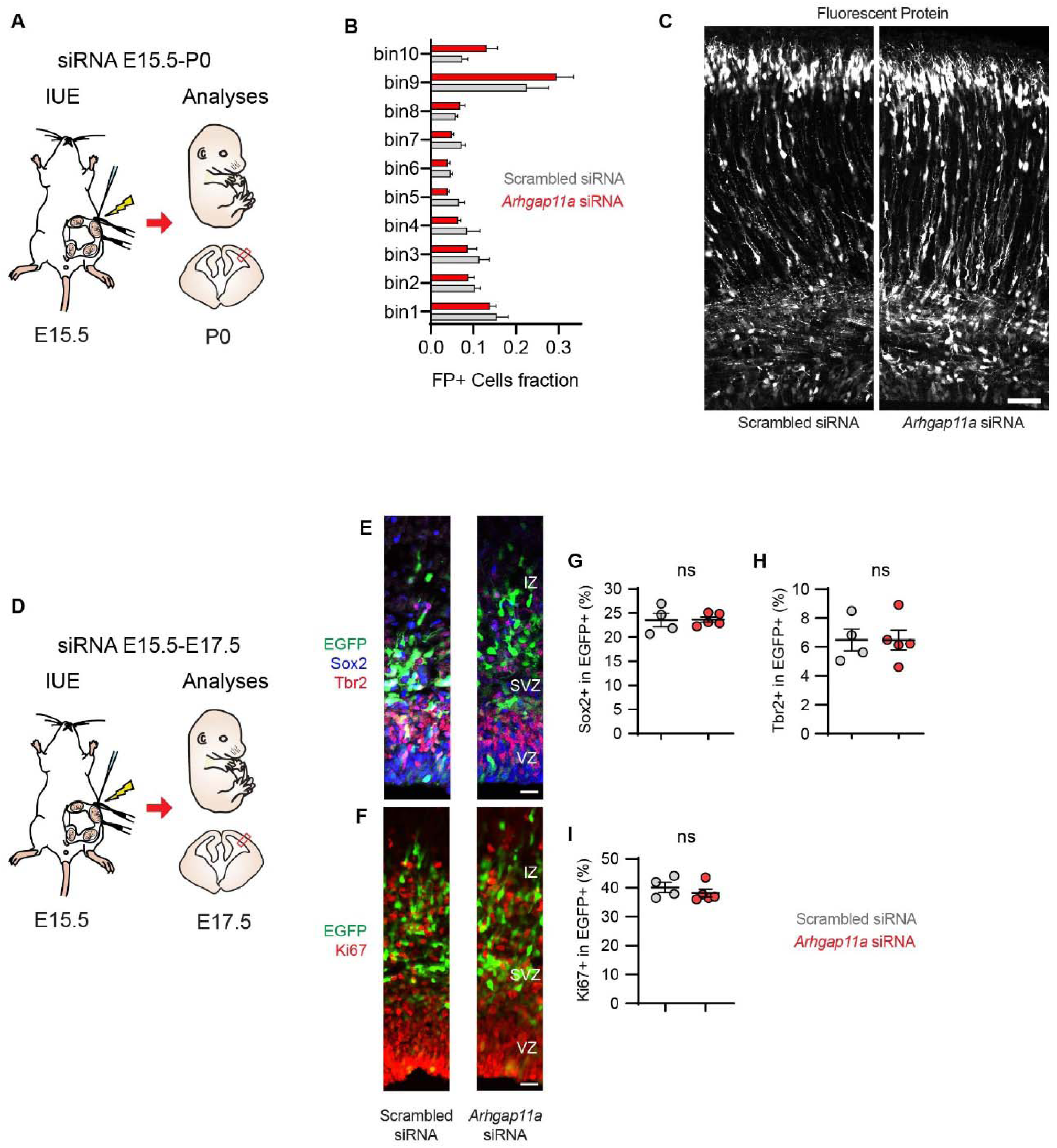
*Arhgap11a* depletion does not impact neural progenitor proliferation and numbers. Related to Figure 3. (A) Schematic overview of experiments to test the cell autonomous impact of *Arhgap11a* depletion on neuron positioning at P0. (B) Quantification of electroporated cell distribution at P0. (Scrambled: n= 3 brains from 2 independent experiments, *Arhgap11a:* n= 4 brains from 2 independent experiments, two-way ANOVA). (C) Representative images of regions analyzed at P0. (D) Schematic overview of experiments to test the impact of *Arhgap11a* depletion in RGCs progenitor proliferation and numbers. (E, F) Representative images of regions analyzed at E17.5. (G,H,I) Quantification of Sox2 (G), Tbr2 (H) and Ki67 (I) following *Arhgap11a* depletion. Note there is no defect in the number of neural progenitor subtypes and proliferation. (Scrambled: n=4 brains from 3 independent experiments, *Arhgap11a*: n=5 brains from 3 independent experiments, unpaired t-tests) IUE: *in utero* electroporation, siRNA: small interfering RNA, FP: fluorescent protein, ns: non-significant, VZ: ventricular zone, SVZ : subventricular zone, IZ: intermediate zone. Scale bars C,E,F: 50μm. Graphs show average values +/-SEM.

**Figure S4.**
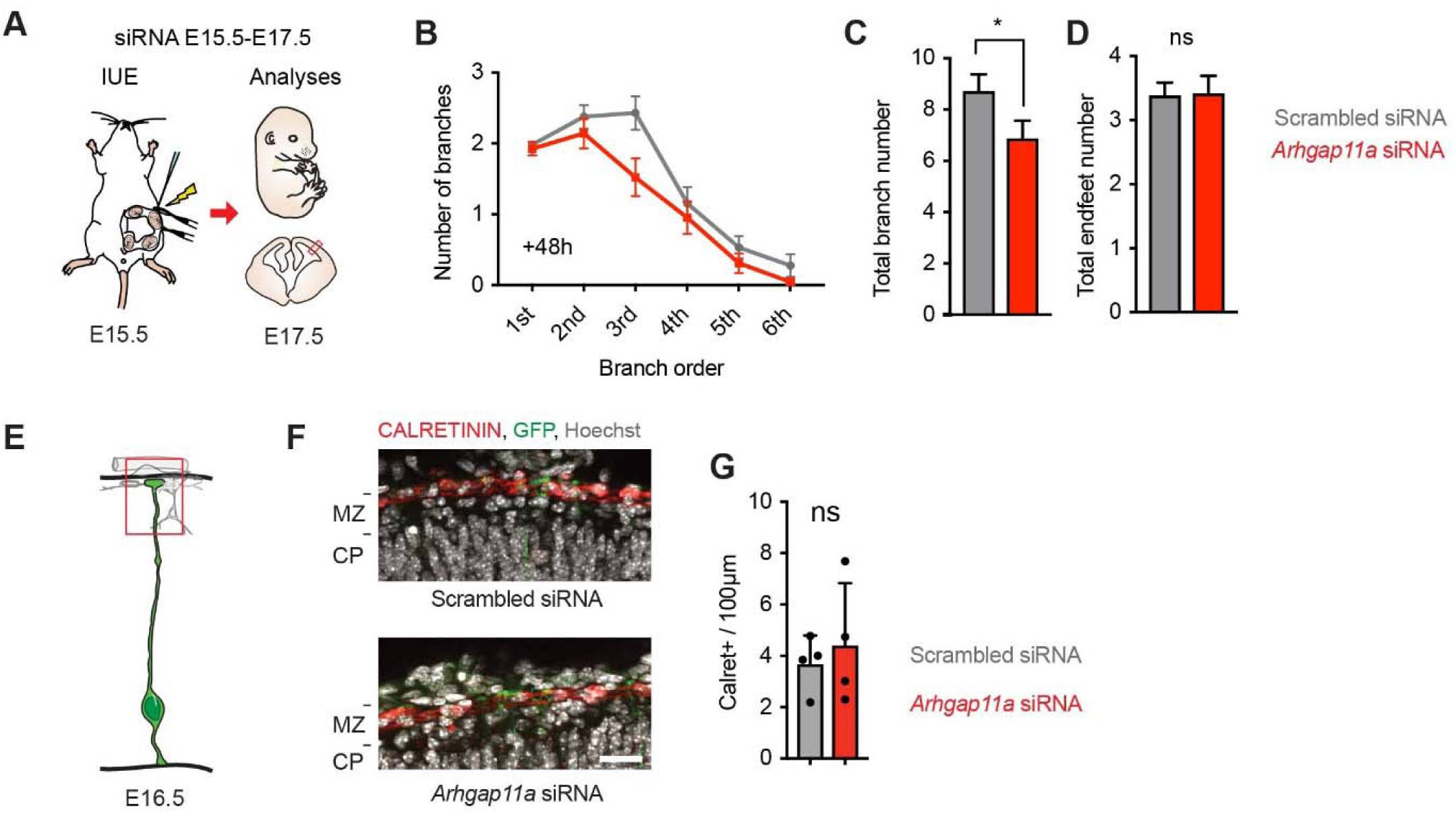
Effect of *Arhgap11a* misexpression in the MZ. Related to Figure 4. (A) Schematic overview of experiments in (B-D) to quantify the impact of acute *Arhgap11a* depletion on basal process complexity two days after electroporation. (B-D) Reduction of RGC basal process complexity persists two days after introduction of *Arhgap11a* siRNAs (B,C) Endfoot number per RGC is not affected by depletion of *Arhgap11a* (D). (Scrambled: n=58 cells from 5 brains, 2 independent experiments, *Arhgap11a:* n=43 cells from 4 brains, 2 independent experiments, C,D: unpaired t-tests). (E-G) A one day knockdown of *Arhgap11a* in RGCs (E) has no impact on the number of CALRETININ+ cells (presumed Cajal Retzius neurons) lining the basement membrane (F, G). (Scrambled: n=4 brains from 3 independent experiments, *Arhgap11a:* n=4 brains from 3 independent experiments, unpaired t-test). IUE: *in utero* electroporation, siRNA: small interfering RNA, *: p-value<0.05, ns: non-significant. Scale bar: F: 25μm.. Graphs show average values +/-SEM.

**Figure S5.**
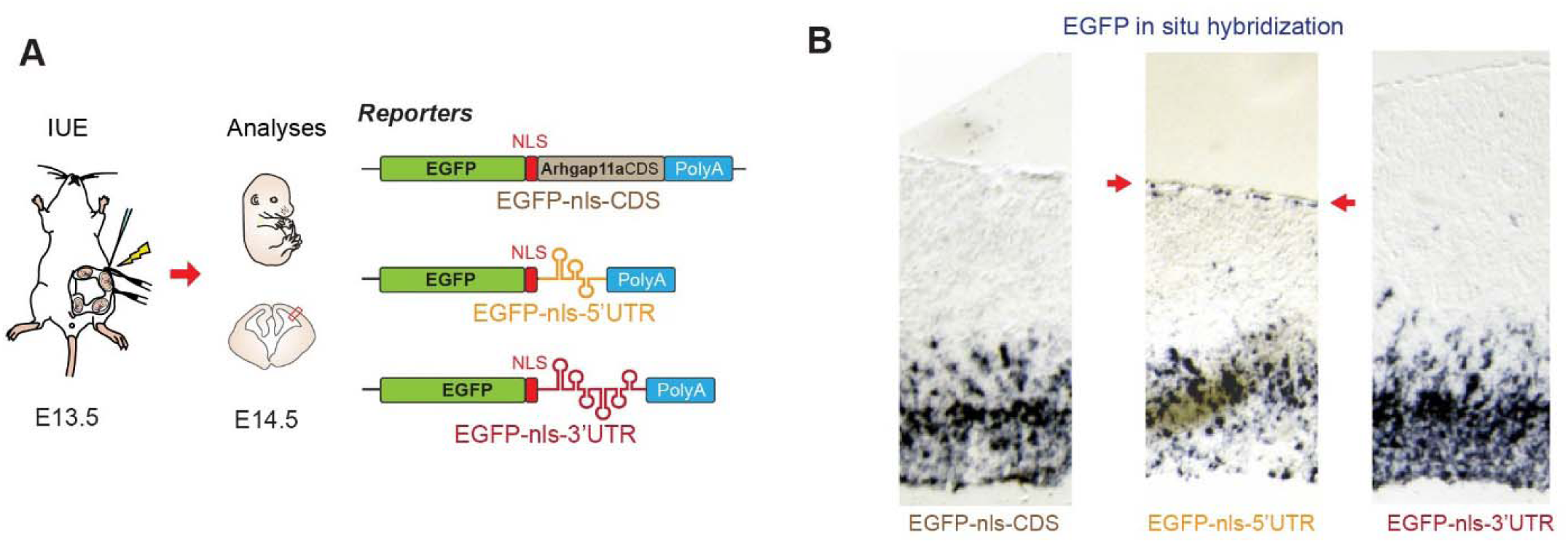
Localization of *Arhgap11a* mRNA to RGC basal endfeet and processes relies on a 5’UTR element. Related to Figure 5. (A) Strategy used to uncover the endfoot localization sequence in *Arhgap11a* mRNA. (B) *In situ* hybridization against *EGFP* (purple) in E14.5 brains. Note EGFP-nls mRNA reporters localize to the pia only when they contain the *Arhgap11a* 5’UTR (red arrows denote enrichment in the basal endfeet region). This shows that *Arhgap11a* mRNA endfoot localization sequence resides in its 5’UTR. IUE: *in utero* electroporation, CDS: coding sequence, UTR: untranslated region, nls: nuclear localization signal.

**Figure S6.**
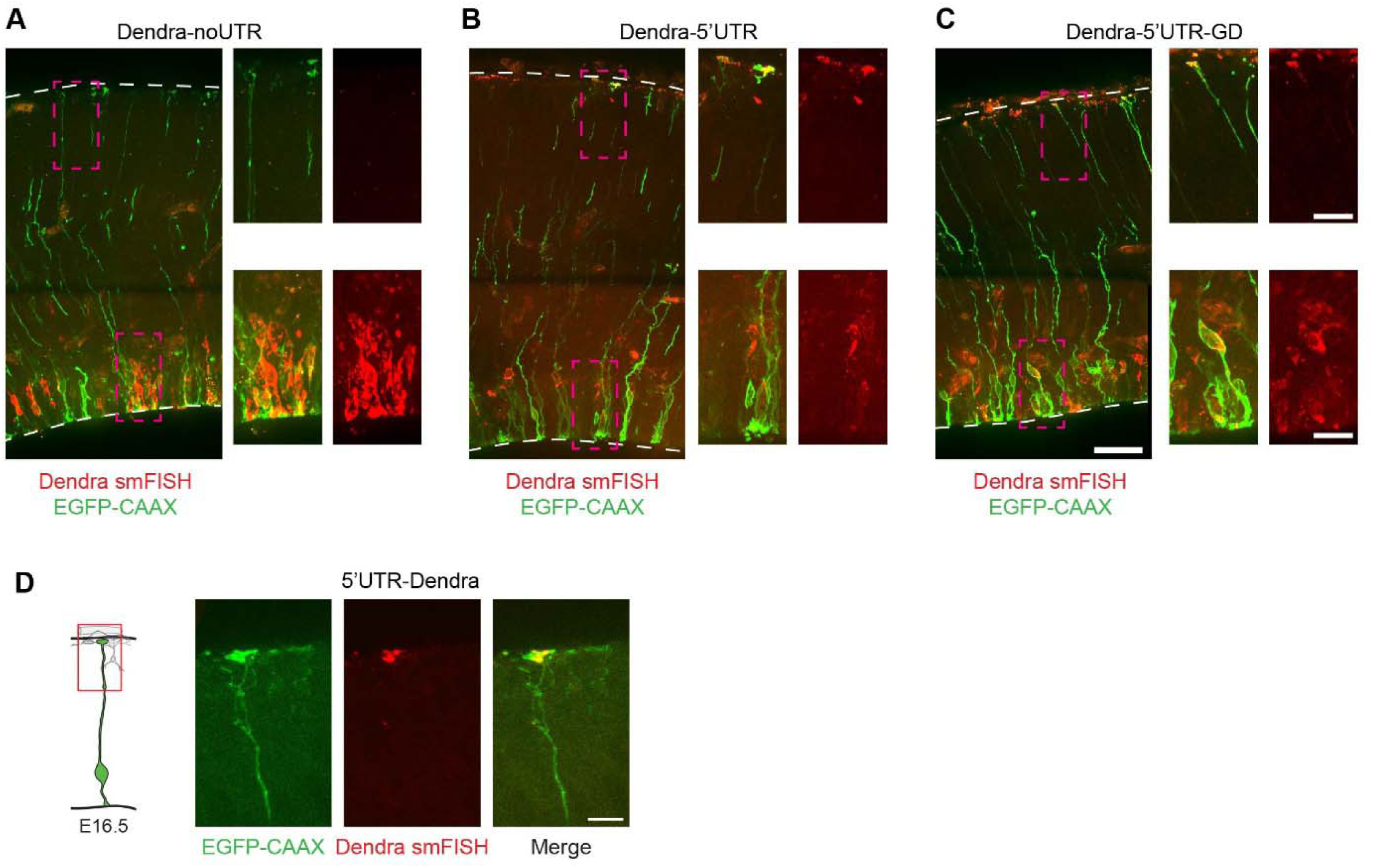
Localization of *Arhgap11a* mRNA to RGC basal endfeet and processes relies on a 5’UTR element. Related to Figure 6. (A) Strategy used to uncover the endfoot localization sequence in *Arhgap11a* mRNA. (B) *In situ* hybridization against *EGFP* (purple) in E14.5 brains. Note EGFP-nls mRNA reporters localize to the pia only when they contain the *Arhgap11a* 5’UTR (red arrows denote enrichment in the basal endfeet region). This shows that *Arhgap11a* mRNA endfoot localization sequence resides in its 5’UTR. IUE: *in utero* electroporation, CDS: coding sequence, UTR: untranslated region, nls: nuclear localization signal.

**Figure S7.**
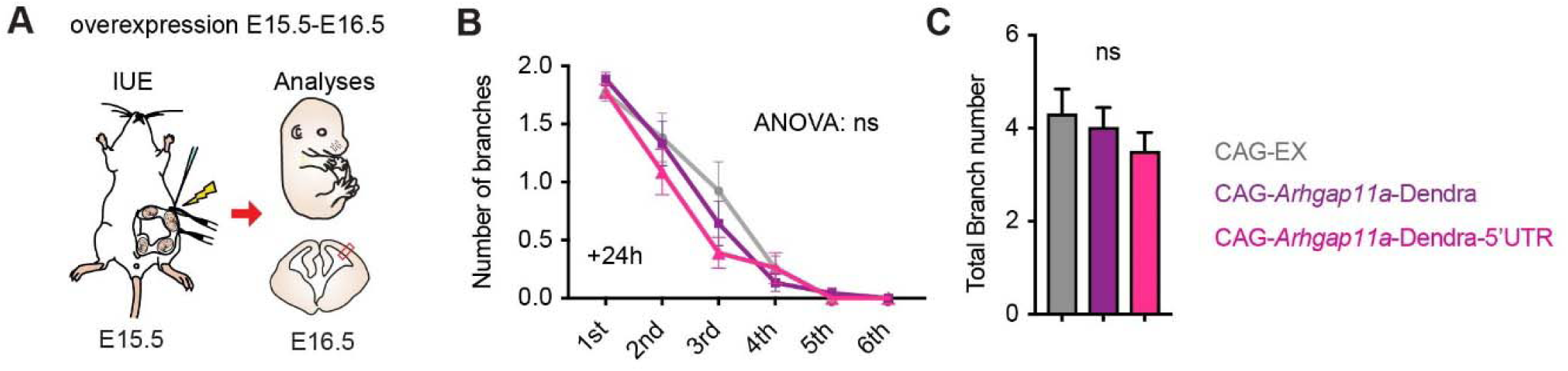
*Arhgap11a* overexpression does not impact RGC basal process complexity in the MZ. Related to Figure 7. (A) Schematic overview of experiments to quantify the impact of acute *Arhgap11a* overexpression on basal process complexity in the MZ, two days after electroporation. (B,C) *Arhgap11a* overexpression does not impact RGC basal process complexity. (CAG-EX: n=39 cells from 3 brains, 2 independent experiments, CAG-*Arhgap11a*-Dendra: n= 36 cells from 3 brains, 2 independent experiment, CAG-Arhgap11a-Dendra-5’UTR: n=45 cells from 4 brains, 2 independent experiments, two-way ANOVA). IUE: *in utero* electroporation, UTR: untranslated region. ns: non-significant. Graphs show average values +/-SEM.

## SUPPLEMENTARY MOVIES LEGENDS

**Supplementary Movie 1**

Time lapse imaging of the upper CP/MZ region of a RGC labeled with membrane-bound EGFP (time: mm:ss).

**Supplementary Movie 2**

Time lapse imaging showing the movement of MS2-labelled mRNAs (green) in radial glial cells within E16.5 mouse organotypic brain slices during one minute. These mRNAs include the *Arhgap11a* cis-regulatory sequence responsible for endfoot RNA localization in radial glia.

